# Hospital and urban wastewaters shape the structure and active resistome of environmental biofilms

**DOI:** 10.1101/2023.01.19.524754

**Authors:** Elena Buelow, Catherine Dauga, Claire Carrion, Hugo Mathé-Hubert, Sophia Achaibou, Margaux Gaschet, Thomas Jové, Olivier Chesneau, Sean P. Kennedy, Marie-Cecile Ploy, Sandra Da Re, Christophe Dagot

**Author notes:** present address: Institut Pasteur, University Paris Cité, Department of Virology, Arboviruses and Insect Vectors, F-75015 Paris, France (Catherine Dauga).

## Abstract

**Background:** Demonstration of the transfer, dynamics, and regulation of antibiotic resistance genes (ARGs) and mobile genetic elements (MGEs) in a complex environmental matrix is yet experimentally challenging, with many essential open questions such as how and where transfer and dissemination of ARGs happens in nature. The extent and conditions of MGEs transfer that carry ARGs is still largely unexplored in natural environments and microbial communities. Biofilms are structures that include high density multi-species bacterial communities embedded in self-produced extracellular polymeric substances (EPS) constituting a matrix that facilitates gene transfer and where bacteria exhibit high tolerance to stress and to antibiotics. In this study we implemented a sampling and analysis approach that allows phenotypic and genomic analyses of *in situ* and reconstituted *in vitro* hospital and urban wastewater (WW) biofilms. To assess the potential of hospital and urban WW biofilms to efficiently disseminate ARGs in the WW system, we explored the EPS within the biofilm matrix and assessed the expression of the resistome (ARGs) and mobilome (MGEs) by metatranscriptomics.

**Results:** We first showed that a) the composition of EPS differs depending on their growth environment (*in situ* and *in vitro*) and their sampling origin (hospital vs urban WW) and that b) a low amount of ciprofloxacin impacted the composition of the EPS. Next, the metatranscriptomic approach showed that a) expression of ARGs and MGEs increase upon adding a low amount of ciprofloxacin for biofilms from hospital WW but not for those from urban WW and b) that expression of specific plasmids that carry individual or multiple ARGs varies depending on the WW origins of the biofilms. When the same plasmids were expressed in both, urban and hospital WW biofilms, they carried and expressed different ARGs.

**Conclusion:** We show that hospital and urban wastewaters shape the structure and active resistome of environmental biofilms, and we confirmed that hospital WW is an important hot spot for the dissemination and selection of AMR. The different responses to antibiotic pressure in hospital *vs* urban biofilms, coupled with differences in biofilm structure helps delineate distinct characteristics of hospital and urban WW biofilms highlighting the relationships between the resistome and its expression in environmental biofilms and their surrounding ecosystems.

## Introduction

Biofilms are the preferred lifestyle of bacteria in the natural environment. They present structures that support high densities of bacteria, exhibit high tolerance to antimicrobials, and facilitate gene transfer [1–3]. In aquatic environments, biofilms are suggested to serve as hot spots for the accumulation and dissemination of antimicrobial resistance genes (ARG) [1,4]. Chemical pollutants spread by urban, industrial, and agricultural waste exert a continuous selective pressure on the diverse antibiotic resistance gene (ARG) pool present in the natural environment and its microbial communities [5–7]. For example, ciprofloxacin is a very stable molecule often found in anthropogenically polluted environments and enriched in wastewater (WW), particularly hospital WW [7–10]. Ciprofloxacin is not efficiently removed by WW treatment through activated sludge [11]; it can be detected in WW and downstream aquatic environments and was identified to pose a significant risk for selection of antimicrobial resistance (AMR) in these environments [9,10,12]. Furthermore, ciprofloxacin is known to induce the SOS-response in Gram-negative bacteria. This induction of the SOS response can lead to the expression of antibiotic resistance genes as the quinolone resistance-encoding *qnrB* gene, or the expression of genetic elements as the class 1 integron integrase gene, leading to ARG cassette rearrangements [13,14]. In addition, bacteria that are present in the human gut carry a wide variety of ARGs that can be spread into the environment by human feces through WW [15–18]. To limit the dissemination of chemical pollutants, pathogens and ARGs into the environment, the implementation of WW treatment systems worldwide plays a key role [6,19–21]. However, WW treatment plants are also regarded as selection and dissemination hot spots due to the high density of human gut, environmental bacteria, and chemical pollutants [22–24]. Biofilms in WW systems represent an important source of ARGs and bacteria that may end up circulating in the environment and hence present an important model to study the dissemination of resistance in the context of discharged effluents [4,25,26]. Biofilms dominate most habitats on earth, accounting for ~ 80% of all bacterial and archaeal cells [27]. Biofilm communities are typified by high densities of bacteria embedded in a self-produced protective matrix, which contributes to high antibiotic tolerance, and facilitates gene transfer [4,28]. Extracellular polymeric substances (EPS), such as external DNA (eDNA), proteins, lipids, and polysaccharides, are essential structural components of the biofilm matrix that are involved in resilience to antibiotics and exogenous stress, but also in horizontal gene transfer (HGT) [29–31]. Previous studies have shown that hospital WW contain significantly higher amounts of resistant bacteria, ARGs and mobile genetic elements (MGEs), as well as chemical and pharmaceutical pollutants compared to urban WW. However, these quantities are generally diluted when hospital WW is mixed with urban WW in community sewer systems [7,32,33].

Demonstration of the transfer, dynamics, and regulation of ARGs and MGEs in a complex environmental matrix is yet experimentally challenging, with many essential open questions such as how and when transfer and dissemination of ARGs occur in nature. The extent and conditions of MGE transfer that carry ARGs is still largely unexplored in natural environments and microbial communities. In this study we implemented a sampling and analysis approach that allows phenotypic and genomic analyses of *in situ* and reconstituted *in vitro* hospital and urban wastewater (WW) biofilms. To better understand the role of hospital and urban WW biofilms in effectively disseminating antimicrobial resistance in the WW system, we explored the extracellular polymeric substances (EPS) within the biofilm matrix, the composition of the biofilm community (microbiota), the resistome (antibiotic, biocide, and heavy metal resistance genes) and mobilome (mobile genetic elements), with focus on expressed genes (metatranscriptome).

## Materials and Methods

### Sample collection and biofilm production *in situ* and *in vitro*

Wastewater (WW) biofilm production campaigns (*in vitro* and *in situ*) were conducted in January 2017. For *in vitro* biofilms, 20L of untreated urban (up flow of the WWTP Limoges city), and hospital (900-bed, Limoges teaching hospital France) WW were collected on the same day and transported (at 4°C) to the laboratory for immediate use in continuous biofilm reactors. The remaining WW was stored at 4°C to replace evaporated volume every two days. For each WW type, two reactors containing 3L of WW were fed at a continuous rate, from an external container containing the same WW (5L in total), ensuring circulation and a stable level in the biofilm reactors (closed continuous system) (Supplementary Figure 1). Rooms were kept darkened, and the temperature was maintained at 17°C. Sixteen polystyrene slides suitable for confocal laser scanning microscopy (CLSM) (Agar Scientific LTD, UK) were attached to a rotator that was constantly turning, producing 16 biological replicate biofilms per reactor. After 3 days of initial biofilm formation, 1μg/L of ciprofloxacin (considered as the minimum selective concentration (MSC) of ciprofloxacin for most environmental bacteria [34] was added to one reactor for each biofilm type (urban, hospital, lab duplicate). On day 7, the urban and hospital biofilms grown in the reactors containing ciprofloxacin were challenged with a fresh dose of ciprofloxacin (1 μg/L) and harvested the same day (D7) 4 hours after ciprofloxacin challenge. Samples were processed within 30 minutes after harvesting. For metagenomics analysis, 8 biological replica biofilms were scraped with tissue cell scrapers (Biologix^®^) from the slides and washed/suspended in 1X phosphate buffered saline solution (PBS), transferred in 2ml Eppendorf tubes, and immediately shock frozen in liquid nitrogen. Samples were stored at −80°C until RNA/DNA extraction. For CLSM analysis, the remaining 8 slides were washed gently once with 1x PBS and either directly stained (for life/dead staining) or fixed in 3.7% formaldehyde solution for 1 hour and preserved for subsequent EPS staining.

For *in situ* biofilms, holes were drilled into polystyrene slides ends and attached with fishing line (16 slides in total, Supplementary Figure 1). The string of polystyrene slides was immersed into the hospital WW pipe and attached at the access point. Slides were recovered after 7 days. Slides were separated and carefully collected in a slide box containing cooled 1xPBS solution and transported to the laboratory immediately for processing. Slides that showed sufficient biofilm formation (by eye) and were least polluted with residues of tissue and other large waste debris, were immediately scraped from the slides and shock frozen in liquid nitrogen as described above. Remaining slides were washed gently in 1xPBS and fixed in 3.7% formaldehyde solution for 1 hour for subsequent EPS staining and CLSM analysis. The same procedure was applied for the *in situ* biofilm production in the untreated urban WW directly at the WWTP (slides were attached and emerged in the large WW receiving pipe/body after primary water filtration step (removal of large debris)). Slides were recovered as described above and stored in a slide box containing cooled 1xPBS. At least four suitable biofilm slides/replicates were directly processed on site by scraping them from the slides and shock freezing them in a portable liquid nitrogen tank. The remaining slides were transported at 4°C to the laboratory and fixed in 3.7% formaldehyde solution for subsequent EPS staining and CLSM analysis as described below.

### Sample processing

DNA extraction was performed with the DNA/RNA dual extraction kit (Qiagen) according to the manufacturer’s instructions and stored at −20°C. RNA quality, as measured by RNA integrity number (RIN) was insufficient for RNA sequencing. Therefore, RNA was extracted from remaining biological replicates and samples using the ZymoBIOMICS DNA/RNA Miniprep Kit. DNA concentration was determined by Qubit Fluorometric Quantitation (Thermo fisher scientific, Waltham, MA USA) assays according to the manufacturer’s instructions. All DNA samples were diluted or concentrated to a final concentration of 10 ng/μl for downstream qPCR and 16S rRNA analysis. RNA concentration and integrity were analyzed by the Agilent 2100 Bioanalyzer system.

### 16S rRNA gene sequencing and sequence data pre-processing

Extracted DNA samples for 16S rRNA sequencing were prepared following a dual barcoded two-step PCR procedure for amplicon sequencing for Illumina. The V4 region was sequenced by generating 2 × 300 bp on an Illumina MiSeq. Data was demultiplexed and filtered for quality and remaining primers according to recommended practices as described previously [7].

### 16S rRNA data analysis

Illumina MiSeq forward and reverse reads were processed using the MASQUE pipeline (https://github.com/aghozlane/masque). Briefly, raw reads are filtered and combined followed by dereplication (dereplication is the identification of unique sequences so that only one copy of each sequence is reported) Chimera removal and clustering are followed by taxonomic annotation of the resulting OTUs by comparison to the SILVA database. A BIOM file is generated that combines both OTU taxonomic assignment and the number of matching reads for each sample. Relative abundance levels for bacterial taxa (family/genus level) were obtained and analyzed [7].

### High-throughput qPCR for the characterization of the resistome composition and normalized abundance

Nanolitre-scale quantitative PCRs to quantify genes that confer resistance to antimicrobials were performed as described previously [7,32,35]. We targeted 76 genes, grouped into 16 resistance gene classes; targeted genes include ARGs that are prevalent in the gut microbiota of healthy individuals [15], clinically relevant ARGs (extended-spectrum β-lactamases (ESBLs), carbapenemases, and vancomycin resistance), and heavy metals and quaternary ammonium compounds resistance genes suggested to favor cross and co–selection for ARGs in the environment [36,37]. We also targeted 11 other targets, including common transposase gene families and class 1, 2 and 3 integron integrase genes, that are important vectors for ARGs in the clinics and often used as proxy for anthropogenic pollution [38]. This set of 87 genes constituted what we named the targeted resistome (Supplementary Table 1). We also included primers targeting 16S rDNA. Primer design and validation prior to and after Biomark analysis has been done as described earlier [7,35]. Real-Time PCR analysis was performed using the 96.96 BioMark™ Dynamic Array for Real-Time PCR (Fluidigm Corporation, San Francisco, CA, U.S.A) as described previously (7). Thermal cycling and real-time imaging were performed at the Plateforme Génomique GeT – INRA Transfert (https://get.genotoul.fr/en/), and Ct values were calculated using the BioMark Real-Time PCR analysis software. Calculations for normalized and cumulative abundance of individual genes and allocated gene classes were performed as previously described [7,32,35]. The resistome data displayed here and used for statistical analysis is based on resistome data that was obtained for two biological replicates per sample.

### Metatranscriptomics and RNA sequencing

Cell pellets were re-suspended in TRIzol reagent and subjected to three cycles of 1min bead-beating (0.5mm silica/zirconia beads) followed by RNA extraction using the Direct-zol RNA Kit (Zymo research) according to the manufactures’ protocol. DNA was removed by TURBO DNase (Ambion) until no genomic DNA was detected by PCR. The quality and integrity of the total RNA were checked on the Bioanalyzer system (Agilent). Because RNA yield is low, only 6 samples containing the highest amount of RNA and best RIM values (>6) were selected for RNA sequencing and meta-transcriptomic analysis (data not shown). Ribosomal RNA depletion was performed using the Bacteria RiboZero kit (Illumina). From rRNA-depleted RNA, directional libraries were prepared using the TruSeq Stranded mRNA Sample preparation kit following the manufacturer’s instructions (Illumina). Libraries were quantified on Bioanalyzer DNA chips (Agilent) and Qubit^®^ dsDNA HS Assay Kit (ThermoFisher). 150 bp paired read sequences were generated on the Nextseq 500 (high output) sequencer according to manufacturer’s instructions (Illumina). The multiplexing level was 10 samples per lane. Reads were cleaned for low-quality and adapter sequences using Cutadapt version 1.11.

The level of ribosomal RNAs (rRNAs) remained high in our environmental samples, even after both *in vitro* rRNA depletion step and optimized RNA extraction methods (see above). We tackled these technical shortcomings by including an *in-silico* rRNA depletion step to improve mRNA read assembly and subsequent analysis (Supplementary Fig. 2 and Supplementary Tables 2 and 3). Ribosomal RNA reads were filtered out by mapping on the Silva database with BWA-MEM v0.1.17 (default parameters) and Samtools v1.9 (36). Then, unmapped reads (mRNA) were subjected to *de novo* assembly using SPADES v3.13.1 (k-mer length 21). From 31% to 42% of trimmed reads coding for rRNA per sample were fast assigned to bacterial families with Kraken [39] against the Silva SSU/LSU-Ref database (Release 132) [40]. 27% to 31% of reads corresponded to not classifiable metagenomics sequences, related to microbial dark matter [41]. 0.68% to 12% of reads were classifiable on the 16S rRNA Silva database. (Supplementary Figure 6, Supplementary Table 3).

After sequencing, trimming and *in silico* ribosomal RNA depletion, we obtained 4.137.706 putative mRNA reads (57.77 % of the initial sample) for the Hospital *in situ* biofilm, and 35.861.826 putative mRNA reads (29.6% of the initial sample) for the Urban *in situ* biofilm, leading to a reasonable number of contigs to analyse, with 866.221 and 2.379.394 contigs for the hospital and urban WW *in situ* biofilms, respectively (Supplementary Table 2). For the hospital WW *in vitro* biofilms HL-1 and HL-2 (HL-2 = hospital WW *in vitro* biofilm with ciprofloxacin added) 14.860.486 and 36.452.020 mRNA reads were obtained leading to 2.332.472 (HL-1) and 3.285.208 (HL-2) assembled contigs, respectively. For the urban WW *in vitro* biofilms UL-1 and UL-2 (UL-2= urban WW *in vitro* biofilm + ciprofloxacin), 55.513.773 and 38.054.494 mRNA reads were obtained leading to 3.483.745 and 2.684.771 assembled contigs, respectively (Supplementary Table 2 and 3).

### Metatranscriptomic analysis

Mapping the mRNA assemblies to specific reference databases enables to assess the expression levels of functionally viable ARGs within biofilms.

A specialized database of the genes targeted by the high-throughput qPCR for the targeted resistome analysis was built (accession numbers available in Supplementary Table 1) and used for mapping assembled contigs with BWA-MEM v0.1.17 (with default parameters) and SAMtools v1.9. Then, the transcriptome activity and abundance of ARGs was assessed by mapping reads on assembled contigs [42]. Reads were mapped against each ARG-like sequence identified from meta-transcriptomic assembly contigs, using Bowtie2 v2.3.4.3 with default settings. The ARG-like sequences with > 100 mapped reads were assigned to ARG transcripts.

To increase the chance of detecting active ARGs in hospital and urban wastewater biofilms, transcripts coding for ARGs were searched against the CARD database (release 08/21/2019) by using the Resistance Gene Identifier (RGI 3.1.0) retaining perfect and strict hits only [43]. The ARG-like sequences were identified as ARG transcripts when their coverage was up to 80% with a minimum of 100 reads mapping. The expression level of a particular ARG transcript in each sample was calculated by summing the number of hits by ARG class (see list of categories in Supplementary table 3). In addition, we weighted the identification with the mean of read coverage by ARG class.

In addition, we detected ARG-carrying MGEs in WW biofilms, namely the active resistance mobilome. First, assembled contigs were mapped against the ACLAME database [44], using BWA and Samtools. Subsequently, identified MGE-carrying contigs were again mapped against the CARD database to identify ARG-carrying MGEs. If one MGE-carrying contig mapped to different ARGs, only hits with the best CIGAR (extracted from the SAM file) were considered.

All steps described above were implanted in python-based modules, and all modules formed the framework of the pipeline AROM, for ARg On Mobilome (Supplementary Figure 2). Lately, we added the module get abundance to normalize gene expression levels according to the sample size (Gb), the length of reads and the length of each ARG [45]. All changes to AROM (https://github.com/sophiaachaibou/ARGmodules) are tracked in GitHub and the versions managed using bioconda.

### Extracellular polymeric substances (EPS) staining for confocal scanning laser microscopy

The following stains were used to fluorescently label EPS components (polysaccharides, lipids, and external DNA) and to assess live-dead ratios of different biofilm samples. Life-dead: Propidium Iodide (red; 0.5μM working solution) and SYTO-BC (green; Life technologies™; 5uM working solution). 500μl of each fluorescent dye applied and allowed to incubate at ambient temperature in the dark for 30 min. Slides were subsequently rinsed three times with phosphate buffered saline (PBS) and immediately analyzed by CSLM. For the EPS components, biofilms were fixed in 3.7% formaldehyde solution (see above) prior to EPS staining. For external DNA (eDNA) the combination of TOTO-1 (green) and SYTO-60 (red) (Life technologies™) was used. Staining was performed as described by Okshevsky et *al*. [46], by using a final working stock solution in PBS of TOTO-1 at 2μM and SYTO-60 at 10μM. Here, following three PBS washes, slides kept at 4°C in the dark until CSLM analysis the same day. For polysaccharides (carbohydrate binding proteins) and proteins a combination of Concanavalin A, tetramethylrhodamine conjugate (ConA; Thermo Fischer Scientific) selectively binding to α-mannopyranosyl and α-glucopyranosyl residues (carbohydrates) and Fluorescein isothiocyanate isomer I (FITC; Siegma Aldrich) selectively binding to proteins was used. FITC working solution was concentrated at 200μg/ml. 500μl of 0.1M sodium bicarbonate buffer was added to the biofilm slide prior to adding 500μl of FITC staining solution (2μg/ml) and incubated at RT in the dark for 30 minutes. Afterwards the excess stain was removed by washing the biofilm slide gently with 1 x PBS solution for three times. Subsequently 500μl of ConA staining solution (50μg/ml) was added to the slide and incubated for 30 minutes at RT in the dark. Excess stain was removed by washing three times with 1x PBS solution. Slides were stored at 4°C until CSLM analysis. For lipid and internal lipid droplet staining Nile Red (0.1μg/ml) was used in combination with FITC (2μg/ml). Incubation and staining were performed as described above by first applying 500μl of 0.1M sodium bicarbonate buffer, subsequently 500μl of FITC working solution, incubation for 30 minutes at RT in the dark, washing with PBS for removal of excess FITC staining and addition of 500μl of Nile Red working solution, incubation for 30 minutes at RT in the dark, removal of excess dye by washing with PBS three times, and storing at 4°C until analysis by CSLM.

### Confocal scanning laser microscopy settings and analysis

Z stacking acquisitions were acquired with a confocal microscope (ZEISS LSM880) using a x20 and x100 magnification. Five Z stacks were performed per condition. The following settings were used to measure the fluorescence from the samples: autofluorescence (λexc 405nm/λem 460nm), FITC (λexc 488nm/λem 525nm), tetramethylrhodamine (λexc 561nm/λem 580nm), Nile Red (λexc 641nm/λem 650nm), Syto60 (λexc 641nm/λem 678nm) and TOTO-1 (λexc 514nm/λem 533nm).

3D view analysis was performed with the Imaris software (Oxford Instrument) using acquisitions with the x20 magnification. Surfaces were created using a manual thresholding. The quantitative analysis was performed with the open-source software BiofilmQ (https://drescherlab.org/data/biofilmQ/) on the acquisitions made with the x100 magnification. Each fluorescent z-stack was segmented using an automatic thresholding (otsu algorithm) allowing object detection and measurements. We focused on the relative abundance that is the measurement of the proportion of the biovolume occupied by each component (i.e one staining). This biovolume must be understood as the total volume occupied by staining(s) of interest. The relative abundance of the polysaccharides (stained by concanavalin A) and the lipids (stained by nile red) were expressed as a percentage of the biovolume resulting from the merge between concanavalin A OR fluorescein, and between nile red OR fluorescein respectively. eDNA relative abundance was expressed as a percentage of the TOTO-1 staining (i.e eDNA) regarding the biovolume resulting from the merge between TOTO-1 OR SYTO-60 stainings.

### Statistical analysis

#### Biofilm matrices

To test for the effect of sample origin (hospital vs urban WW), environment (*in situ* vs *in vitro*) and addition of ciprofloxacin on the composition of the biofilm matrices, the relative abundance of three features were measured, the eDNA, lipid and polysaccharides. For each of these features three models were built. The first type of model focused on data obtained without ciprofloxacin and analyzed for each measured feature, the effect of the variables ‘sample origin’ and ‘environment’, as well as their interactions. The second type of model focused on data obtained in the *in vitro* environment and analyzed for each of the three measured features, the effect of the variables ‘sample origin’ and ‘ciprofloxacin’, as well as their interactions. The last type of model is devised to perform a pairwise comparison of each sample, hence, a different model was built for each pair of samples, with one explanatory variable and two levels, corresponding to the sample ID. All these tests were based of the Randomization of Residuals in a Permutation Procedure implemented by the R package RRPP.

#### Pairwise comparison of the normalized abundance of the resistome data using the Tukey HSD test

To compare the relative abundances of genes classes between samples, for each class and biological replicate, we summed the normalized abundances of individual genes per class. Then for each gene class, we fitted the relative abundance with a log linear model with one explanatory variable, and one level for each sample. Finally, we applied pairwise comparisons of samples using the Tukey HSD test implemented by the multcomp R package and p-values were adjusted using the single step method [47].

#### Correlation of the normalized abundance for ARGs detected by qPCR and their likelihood to be expressed and detected by our meta-transcriptomic pipeline

To detect the threshold for normalized abundance values of genes measured by qPCR, below which meta-transcriptomic has a low chance of detecting the expression of the gene, we fitted a binomial generalized linear model (GLM) explaining the presence or absence of mapped reads (obtained for individual genes detected by our meta-transcriptomic pipeline) by the normalized abundance of the respective genes measured by qPCR. This allowed to obtain a cut-off value for normalized abundance determined by qPCR that is predicted to have a chance of being expressed and detected by our meta-transcriptomic pipeline. We also fitted a Poisson GLMM to the exact read maps for genes detected by our meta-transcriptomic pipeline, with the same explanatory variable as in the binomial GLM, and in addition, an observation level random effect, to account for overdispersion [48].

#### Pairwise comparison of biofilm communities, mobilome and resistome

For the six types of multivariate description of the biofilms, communities (family and genus levels), mobilome (plasmid and gene levels) and resistome (targeted and non-targeted), we compared each pair of samples by measuring the proportion of the Simpson’s diversity that was between samples. Simpson’s diversity measures the probability that two entities taken at random from the dataset are of the same type. It is calculated as 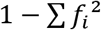, where values of *f* are the frequencies of the entities. To compute the proportion of this Simpson’s diversity that is between a given pair of samples A and B, we computed the total diversity using the average of the frequencies of the two samples and the average diversity within the two sample:

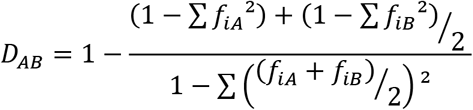

This divergence index turns out to also be the formula of the between population fixation index (*F_ST_*) used in population genetics, using alleles frequencies [49].

Based on these pairwise dissimilarities, we built UPGMA trees (unweighted pair group method with arithmetic mean). The stability of the trees was assessed by bootstrapping the individual genes and taxa. To assess the significance of the divergence between samples we used the pair of samples with the lowest divergence as a conservative reference. Specifically, we categorized bootstraps according to the pair of samples with the lowest divergence, and then calculated the proportion of each category. Finally, we assessed and tested the correlation between the dissimilarity matrices used to build the UPGMA trees using some Mantel tests (R package ade4).

#### Multivariate analysis for ARG detected on plasmids (mobilome)

To analyze the association between the class of the ARGs of the mobilome and the WW origin of the biofilm and their treatment (*in situ, in vitro*, with or without ciprofloxacin), we first performed a correspondence analysis (CA; function ‘dudi.coa’ of the ade4 R package) analyzing the link between the relative abundance of each ARG associated to a MGE. The significance of this association was tested through a χ^2^-test with a p-value based on 10^5^ Monte Carlo simulations, and its effect size through the Cramér’s V statistic, a measure of the association between two categorical variables that varies from 0 (no association) to 1 (complete association). Then ARGs were grouped into gene classes, and the association between gene classes and biofilm origin and treatment was investigated by applying between class analysis (function ‘bca’ of the ade4 R package) to the CA. We refer to this overall analysis as to a between correspondence analysis (bCA) which significance was assessed though a 10^5^ permutation test. This bCA recovered 33% of the total variation.

## Results

### Effects of minimal selective concentrations (MSC) of ciprofloxacin, sample origin and environment, on hospital and urban WW biofilms’ extracellular polymeric substances (EPS)

We first analyzed the global structure of the hospital and urban WW biofilms. The EPS (eDNA, lipids and polysaccharides) were visualized by confocal scanning laser microscopy (CSLM). 3-D imaging (IMARIS) and quantitative analysis (Biofilm Q) showed that the EPS significantly differ between their growth environment, namely whether biofilms were grown *in situ* or, as reconstituted biofilms, *in vitro*. We detected different relative abundance of eDNA (in both hospital and urban biofilms compared to their *in vitro* counterparts), polysaccharides (for urban *in situ* vs *in vitro* + cip biofilms) and lipids (for hospital *in situ* vs *in vitro* biofilms) (Figure 1 and 2, Supplementary Figure 4a and 4b, Supplementary Table 5). We also detected significant differences in the relative abundance of EPS components dependent on the sample origin, meaning between hospital and urban *in situ* WW biofilms, for eDNA and lipids (Figures 1 and 2, Supplementary Figure 4a and 4b). The relative abundance of eDNA was higher for the hospital *in vitro* biofilms compared to the urban *in vitro* WW biofilms. We also assessed the impact of adding ciprofloxacin on the life-dead ratio and EPS components of the *in vitro* reconstituted hospital and urban WW biofilms. A significant increase of dead cells was detected for the hospital *in vitro* biofilm after adding ciprofloxacin, compared to its control (Supplementary Figure 3). The relative abundance of eDNA was significantly lower in the hospital *in vitro* biofilm exposed to ciprofloxacin and significantly lower for polysaccharides in the urban *in vitro* biofilm exposed to ciprofloxacin compared to their control.

**Figure 1:**
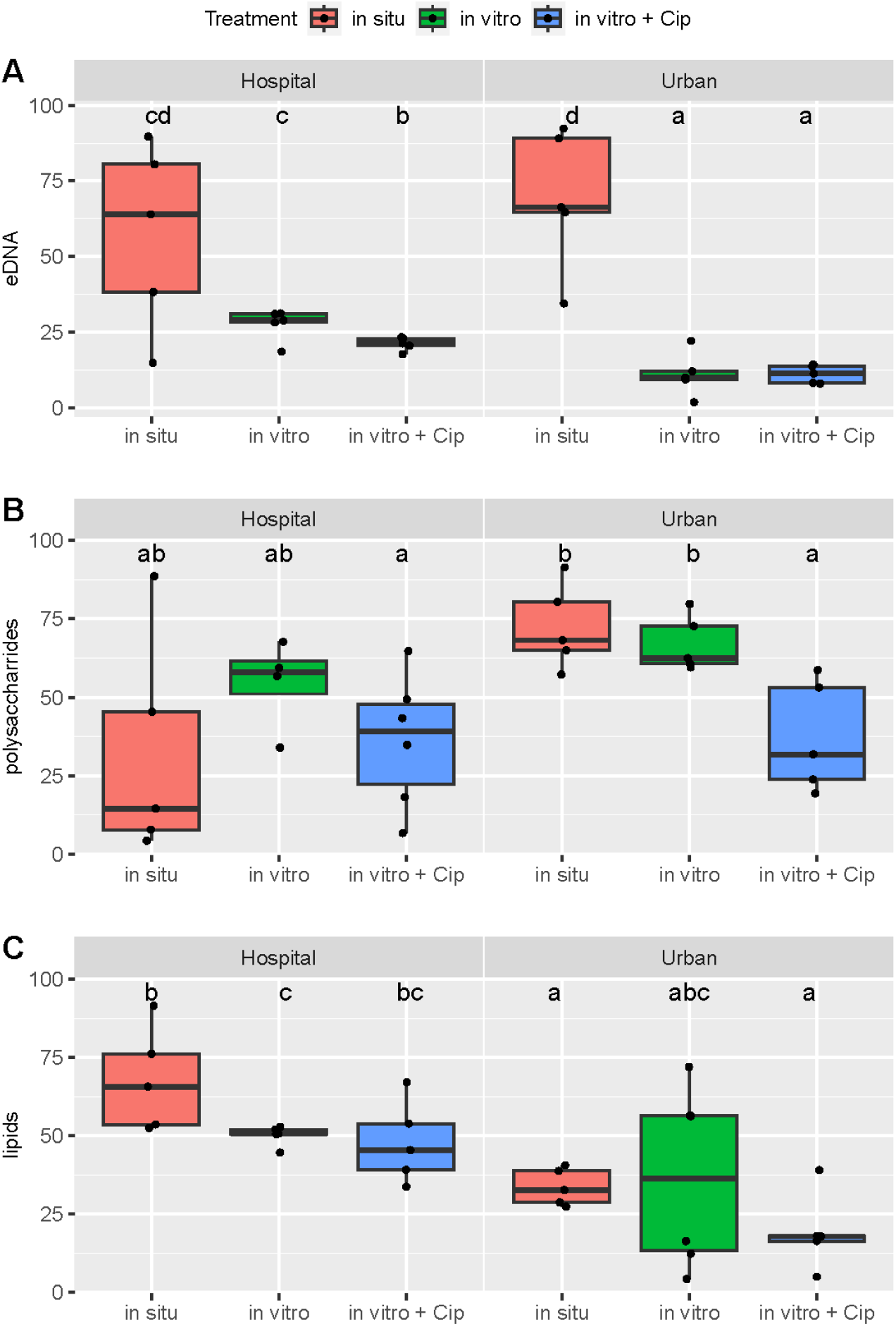
Relative abundance of eDNA (A), polysaccharides (B) and lipids (C) in the respective biofilms. Each EPS component was stained in a different biological replicate, and fluorescence was detected by 5 z-stack acquisitions for each biofilm sample with a 100-fold magnification for the respective biofilms. Quantitative image analysis was performed with Biofilm Q. Results of pairwise permutations tests are summarized using the compact letter display. Samples that do not share a letter in common are significantly different from each other. +Cip= *in vitro* biofilms were exposed after 3 days of initial biofilm formation with 1 μg/ml ciprofloxacin for 4 consecutive days and harvested after exposure of 4 hours on the final day.

**Figure 2:**
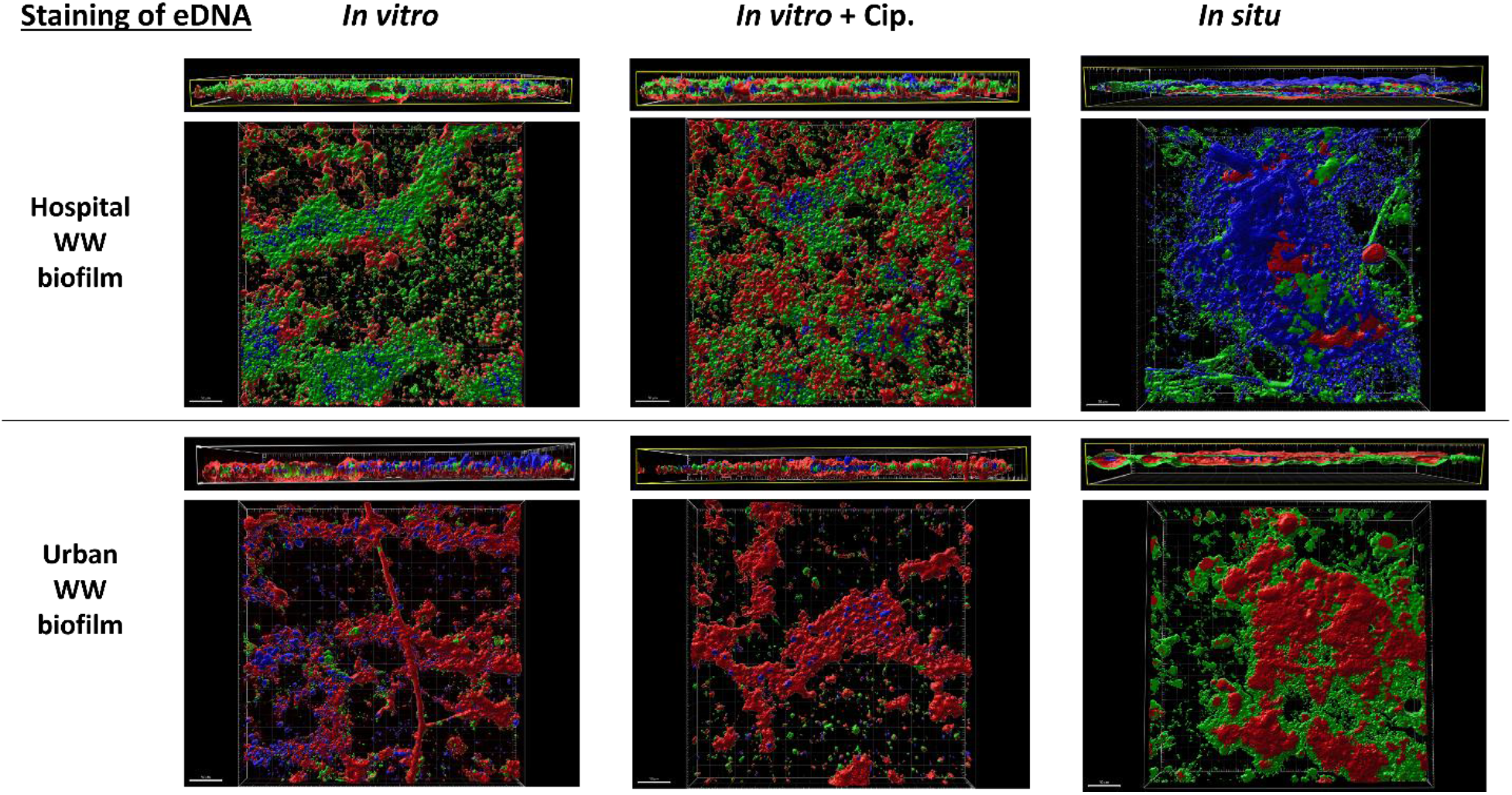
3D imaging of eDNA (green) and total nucleic acids (red), for hospital and urban *in-situ* and *in-vitro* biofilms. Autofluorescence for unknown compounds is in blue. +Cip= *in vitro* biofilms were exposed after 3 days of initial biofilm formation with 1 μg/ml ciprofloxacin for 4 consecutive days and harvested after exposure of 4 hours on the final day.

Finally, we implemented statistical models based on random permutations testing for an overall effect of the environment (*in situ* vs *in vitro*), the sample origin (hospital vs urban) and the effect of ciprofloxacin, on the EPS components of these biofilms (Supplementary Table 5). We confirmed that i) the sample origin (type of hospital vs urban WW) has a significant impact on the abundance of polysaccharides and lipids, and eDNA ii) whether biofilms are grown *in situ* or *in vitro* has a significant impact on the relative abundance of eDNA and iii) ciprofloxacin has a significant impact on the abundance of polysaccharides for both, hospital, and urban *in vitro* biofilms (Supplementary Table 5).

### The microbiota of hospital and urban WW *in situ* and *in vitro* biofilms

The microbial compositions of WW biofilms were assessed using 16S rDNA sequencing. Strikingly and in line with the phenotypic analysis of the biofilm structures, bacterial community profiles of hospital and urban *in situ* biofilms were significantly different from their *in vitro* reconstituted counterparts (Supplementary Figure 5 and Supplementary Table 6). The main genera in the *in situ* hospital WW biofilms are Gram-negative pathogens such as *Pseudomonas, Acinetobacter* and *Aeromonas*, responsible for infections in hospitalized patients [50–52] (Figure 3). *In situ* urban WW biofilms are dominated by different genera than hospital WW biofilms, namely *Flavobacterium* and *Arcobacter* but share *Acinetobacter* with hospital *in situ* biofilms. The genus *Arcobacter* is a common member of human WW systems and can rarely act as animal and human pathogen [53,54]. It is more abundant in urban WW biofilms, compared to the hospital WW environment. *Pseudomonas* is a dominant genus in both urban and hospital *in vitro* biofilms. Surprisingly, *Acinetobacter* decreased in the *in vitro* biofilms compared to their *in situ* counterparts. The addition of ciprofloxacin to the WW in our *in vitro* model did not change the proportional microbiota composition for biofilms from urban WW. However, it had a weak but significant effect on the microbiota composition of biofilms from hospital WW (percentage of the Simpson’s diversity between this pair of samples < 0.2%; Figure 3; Supplementary Figure 5). This suggests that exposure of already formed WW biofilms with ciprofloxacin at MSC in our *in vitro* model has no effect, or only a minor effect, on the primary bacterial ecosystems.

**Figure 3:**
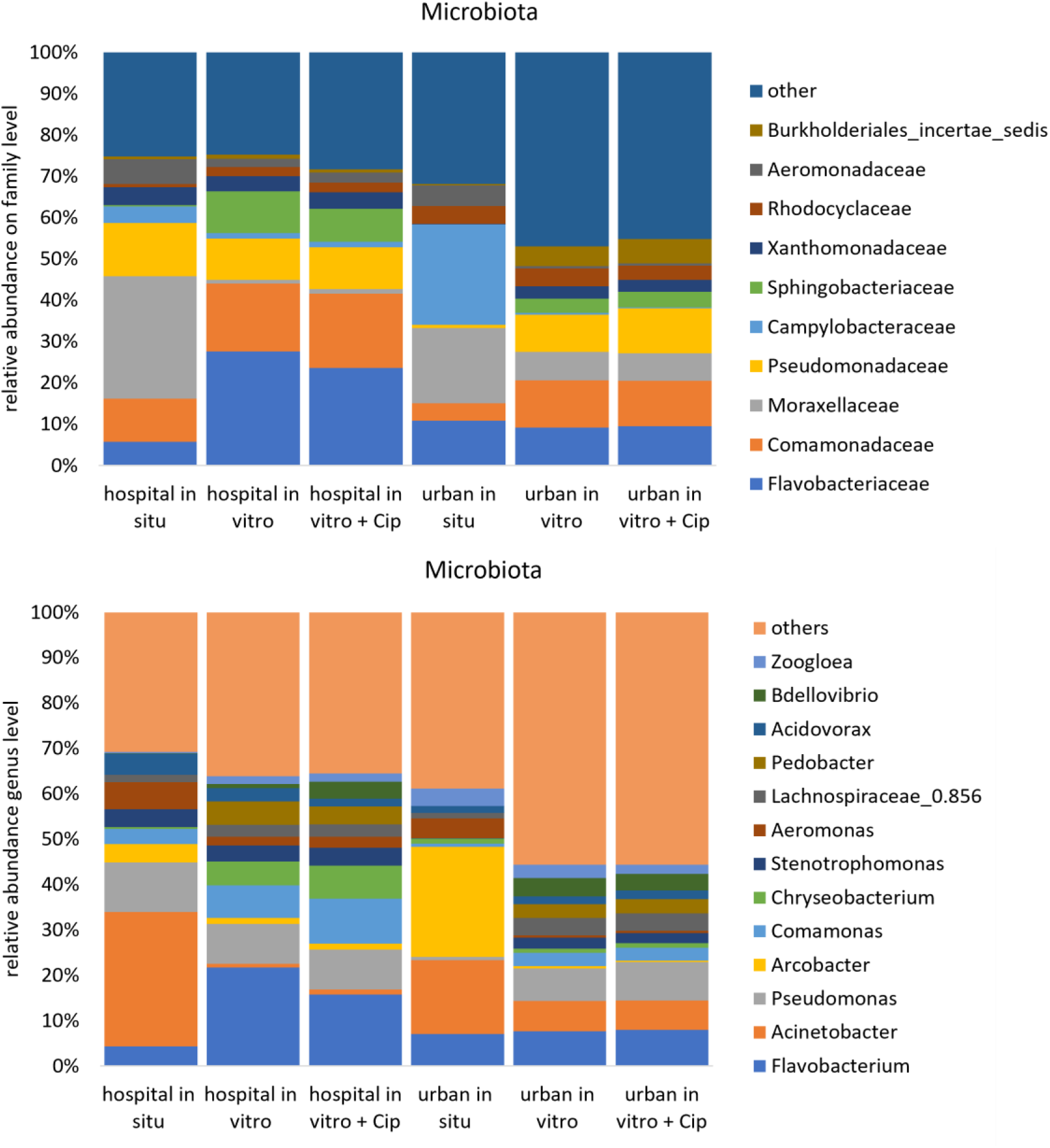
Microbiota composition of hospital and urban wastewater *in situ* and *in vitro* biofilms based on sequencing of the 16S rRNA gene. Depicted are the top 10 taxa on order level and the top 12 on the genus level, the remaining taxa are grouped into others. +Cip= *in vitro* biofilms were exposed after 3 days of initial biofilm formation with 1μg/ml ciprofloxacin for 4 consecutive days and harvested after exposure of 4 hours on the final day (D7).

As described in methods, a relatively large proportion of the RNA reads from the metatranscriptomic RNA sequencing approach of the biofilm communities were ribosomal RNA (rRNA) reads. The 16S rRNA reads were filtered out by mapping on the Silva database, allowing for the assignment of a small proportion of those reads (between 0.68% and 12%) on family level. Hence, this step allowed for additional analysis of the “active microbiota” of these biofilms. Among the classifiable 16S rRNA reads from the metatranscriptomic analysis (methods), we identified a large majority of *Enterobacteriaceae* in hospital biofilms and a mix of *Flavobacteriaceae*, *Enterobacteriaceae*, *Streptococcaceae* and *Francisellaceae* in urban *in situ* biofilm (Supplementary Figure 6 and Supplementary Table 3).

### The targeted resistome and its expression in hospital and urban WW biofilms

To assess the resistome in our samples, a targeted high-throughput qPCR approach, that detects and quantifies 87 individual genes conferring resistance to clinically relevant antibiotics, quaternary ammonium compounds (QACs), heavy metals, transposase genes (MGEs) and integron integrase genes, also referred to as targeted resistome, was applied. At first, comparing the different environments in which biofilms grew (*in situ* vs *in vitro*), we found that for the hospital WW biofilms, all gene classes had a higher normalized abundance in *in situ* biofilm compared to the *in vitro* biofilms, except for aminoglycosides, quinolones, QACs, sulphonamides, MGEs and integrons, which were not significantly different (Figure 4 a and c, Supplementary Figure 7, and Supplementary Table 7). Performing the same comparison for the urban WW *in situ* vs the *in* vitro biofilms, the normalized abundances of chloramphenicol, macrolides, quinolones and MGEs, were significantly higher for the *in situ* biofilms, whereas genes conferring resistance to heavy metals, vancomycin, and sulphonamides had significantly lower normalized abundances in *in situ* urban WW biofilms. Then, comparing the different sampling origin (hospital vs urban WW), the normalized abundance of the targeted resistome was highest in the *in situ* hospital WW biofilms for all detected gene classes apart from macrolides, which were higher in the urban *in situ* biofilm, while tetracyclines, quinolones and MGEs, were not significantly different in their normalized abundance between the two *in situ* biofilm sampling origins (Supplementary Figure 7, and Supplementary Table 7). Comparing the normalized abundance of detected gene classes for the hospital *in vitro* biofilms vs the urban *in vitro* biofilms, it was higher in the hospital *in vitro* biofilms for most genes, except for macrolides, vancomycin and genes encoding for multi-drug efflux pumps for which the normalized abundance was higher in the urban *in vitro* biofilms; no significant difference was observed for tetracyclines and heavy metal resistance genes.

**Figure 4:**
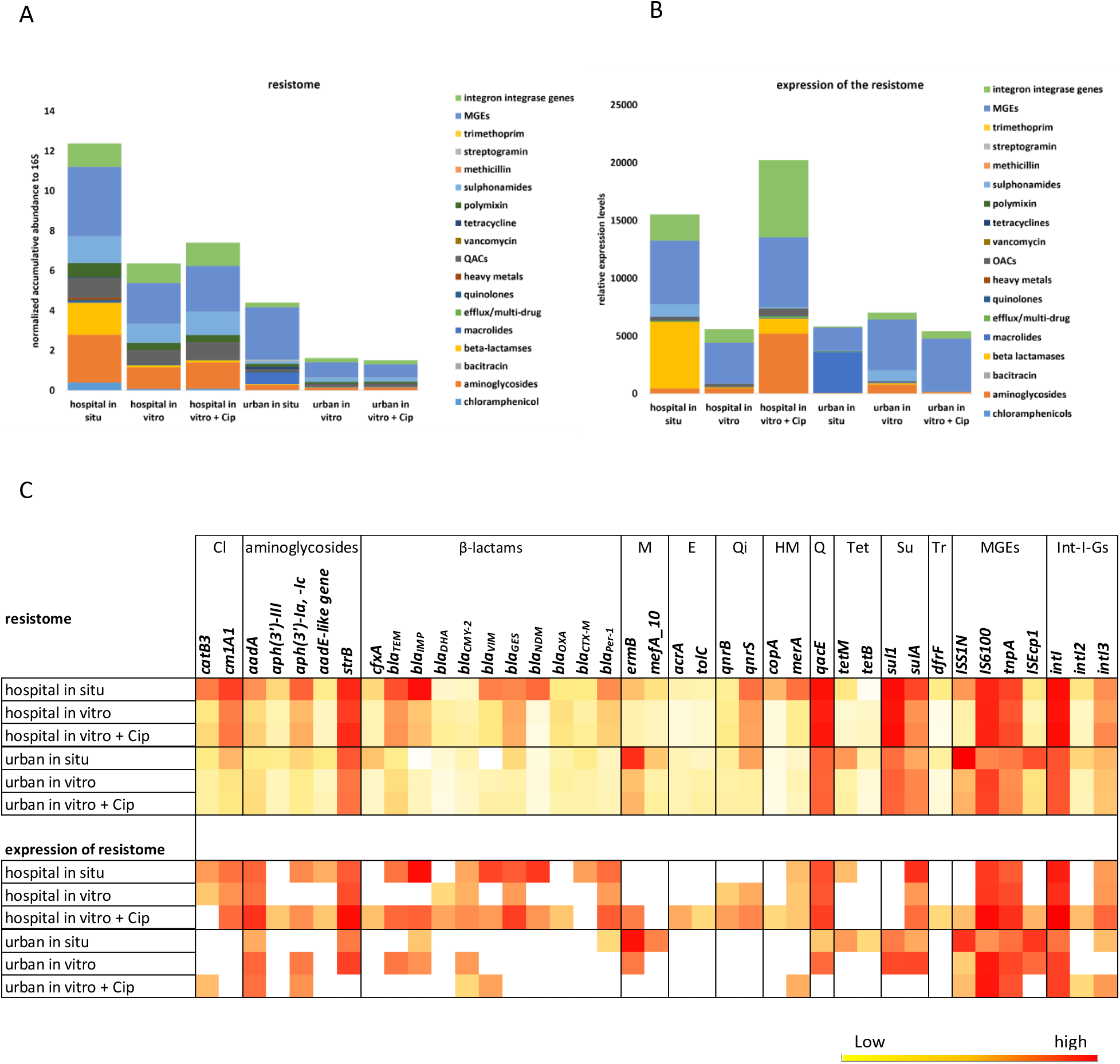
The targeted resistome and its expression in hospital and urban WW biofilms. On panel **A** and **B**, normalized abundance of the targeted resistome (A) and their expression (B) were grouped into gene classes and mobile genetic elements (MGEs) and integron integrase genes. **C**: Heatmap depicting the log2 of normalized abundance of individual genes (targeted resistome) and their normalized expression levels (exact counts of mapped reads for the respective genes) for the individual genes detected by metatranscriptomics. Data were log2 transformed for visualization purpose. +Ab= *in vitro* biofilms were exposed after 3 days of initial biofilm formation with 1μg/ml of the antibiotic ciprofloxacin for 4 consecutive days and harvested after exposure of 4 hours on the final day (D7). Cl= chloramphenicol; M= macrolides; E= efflux pumps/multi-drug resistance genes; Qi= quinolones; HM= heavy metals; Q= quaternary ammonium compounds (QACs); Tet= tetracyclines; MGEs= Mobile Genetic Elements. Int-I-Gs= integron integrase genes class I, II and III. +Cip= *in vitro* biofilms were exposed after 3 days of initial biofilm formation with 1 μg/ml ciprofloxacin for 4 consecutive days and harvested after exposure of 4 hours on the final day.

Then, we analysed the metagenomic messenger RNA (mRNA) to provide information about the targeted resistome actively being expressed in the *in situ* and *in vitro* hospital and urban WW biofilms. Overall, 39 of the 87 targeted genes by qPCR were expressed in at least one of the samples (Figure 4b and c). Following the same comparison pattern, when first comparing the biofilm environments (*in situ* vs *in vitro*), beta-lactam, chloramphenicol and sulphonamide resistance genes were 112, 5, and 47-fold higher expressed in the hospital *in situ* biofilm vs *in vitro* biofilm, whereas macrolide resistance genes were 49-fold higher expressed in the urban *in situ* biofilm compared to *in vitro* biofilm, while for all other gene classes were gene expression could be detected, these detection levels were lower in the urban *in situ* biofilm vs *in vitro* (Figure 4b and c, Supplementary Table 8 and Supplementary Figure 8). For the different sample origins (hospital vs urban WW) of the studied biofilms, expression levels were higher for 7 gene classes, from 5.5-fold higher for genes conferring resistance to aminoglycosides to 957-fold higher for genes conferring resistance to beta-lactams, (Supplementary Table 8 and Supplementary Figure 8) for the *in situ* hospital vs urban *in situ* WW biofilms.

We studied the impact of ciprofloxacin in the *in vitro* biofilm models. When looking at normalized abundance of the targeted resistome in hospital and urban *in vitro* biofilms exposed to ciprofloxacin, no differences could be detected, except for genes conferring resistance to macrolides in the hospital *in vitro* biofilm exposed to ciprofloxacin compared to its control. (Supplementary Figure 7, and Supplementary Table 7). Interestingly, we observed an enhanced expression of the targeted resistome (Figure 4b and c, Supplementary Table 8 and Supplementary Figure 8) after ciprofloxacin addition in the hospital biofilm. For example, expression of ARGs conferring resistance to beta-lactams was 26-fold higher and aminoglycoside resistance genes were expressed 11-fold higher and represented the class of ARGs that was most abundantly expressed proportionally within the hospital biofilm (Figure 4b and c, Supplementary Table 8 and Supplementary Figure 8). Furthermore, ciprofloxacin enhanced expression (4-fold) of the quinolone-resistance genes, *qnrB* and *qnrS* genes, and integron integrase genes (*IntI1* and *IntI3*) by 6-fold and the grouped transposase genes by 1.7-fold compared to its control (Figure 4b and c, Supplementary Table 8 and Supplementary Figure 8). When looking at the detected expression levels of ARG classes in the urban *in vitro* biofilm exposed to ciprofloxacin, expression of ARGs was lower for all detected gene classes apart from the integron integrase and aminoglycoside genes, were expression slightly increased by 1.1 and 1.7-fold respectively, compared to its control (Figure 4b and c, Supplementary Table 8 and Supplementary Figure 8). The *ISS1N* gene was specifically expressed in the *in situ* urban WW biofilm.

Overall, these findings showed that 45% of the targeted resistome is expressed in the WW biofilms and that the expression of ARGs and integron integrase genes in the *in vitro* hospital WW biofilm is enhanced by the addition of ciprofloxacin. Interestingly, genes that were the most abundant in the targeted resistome (qPCR) had a higher probability of being expressed and detected by metatranscriptomics. For example at a threshold of 0.057 for the normalized abundance of genes in the targeted resistome the probability of being expressed and detected by metatranscriptomics is 0.5 (Rho=0.77; Supplementary Figure 10 and Supplementary Table 10).

Below this threshold, the detection of ARG expression could no longer be achieved, suggesting a lack of transcripts. However, technical limitations such as the low sensitivity of meta-transcriptomics and/or the detection of DNA from dead cells and eDNA by qPCR might also be possible explanations as to why not more of the targeted genes could be detected by the metatranscriptomic approach [55–57].

### Assessing the expression of ARGs based on the CARD database

To identify additional ARGs that were not included in our targeted resistome approach, mRNA reads of our samples were mapped against the Comprehensive Antibiotic Resistance Database (CARD) that archives thousands of gene sequences encoding resistance determinants to antimicrobial agents.

ARG expression was higher for *in situ* hospital WW biofilms compared to their *in vitro* counterpart (10.4% for hospital *in situ* biofilm vs about 0.5% for *in vitro* of total mRNA reads in the respective samples (Supplementary Table 1)). The detected expression levels of ARGs in the *in situ* urban WW biofilms was lower compared to their *in vitro* counterpart (0.04% of mRNA reads for the urban *in situ* vs urban *in vitro* 0.4% biofilm). A tenfold higher expression level of ARGs was detected in the *in situ* hospital WW biofilm (10.4 % of all mRNA reads coding for ARGs) compared to the urban WW *in situ* biofilm (0.04 % of mRNA reads coding for ARGs) (Supplementary Table1). For the *in vitro* hospital biofilm exposed to ciprofloxacin, expression level of ARGs was about 4.4-fold higher compared to the non-exposed one (2.4% vs 0.55% vs, (Supplementary Table 1)). No difference in ARG expression level based on total ARG mRNA reads could be observed between urban *in vitro* biofilms exposed or not to ciprofloxacin for the normalized data (0.41% vs 0.37% of total mRNA reads mapping to ARGs, respectively, Supplementary Table 1).

In total we detected the expression of 70 individual resistance genes conferring resistance to 16 classes of antibacterial drugs across all biofilm samples (Figure 5, Supplementary Tables 4 and 9). This approach led to the detection of the expression of additional genes in hospital and urban WW biofilms that were not detected by the targeted approach (Supplementary Table 4). For example, active beta-lactamase such as *bla*^RCP^, *bla*^LCR^, *bla*^FOX^, *bla*^NPS^, four aminoglycoside-modifying enzymes (*aac(3)-Ib/aac(6’)-Ib, aadB, aph(3’’), aph(6)*) and a macrolide phosphotransferase (MPH) were identified. Genes that confer resistance to the antimicrobial classes fusidic acid, elfamycin, rifamycin and antitumoral glycopeptides were found to be actively expressed in all the investigated samples. Genes that confer resistance to isoniazid and triclosan were found to be expressed in the *in vitro* hospital biofilm exposed to ciprofloxacin, and in *in vitro* urban WW biofilms in both conditions. The expression of an enterobacterial fosfomycin resistance gene (*ptsI*) was detected in the *in vitro* hospital WW biofilm exposed to ciprofloxacin (Supplementary Table 4). Furthermore, the expression of multidrug efflux pumps, specifically RND and MFS types, and the expression of the *bla*^OXA^ beta-lactamases was detected in all the samples.

**Figure 5:**
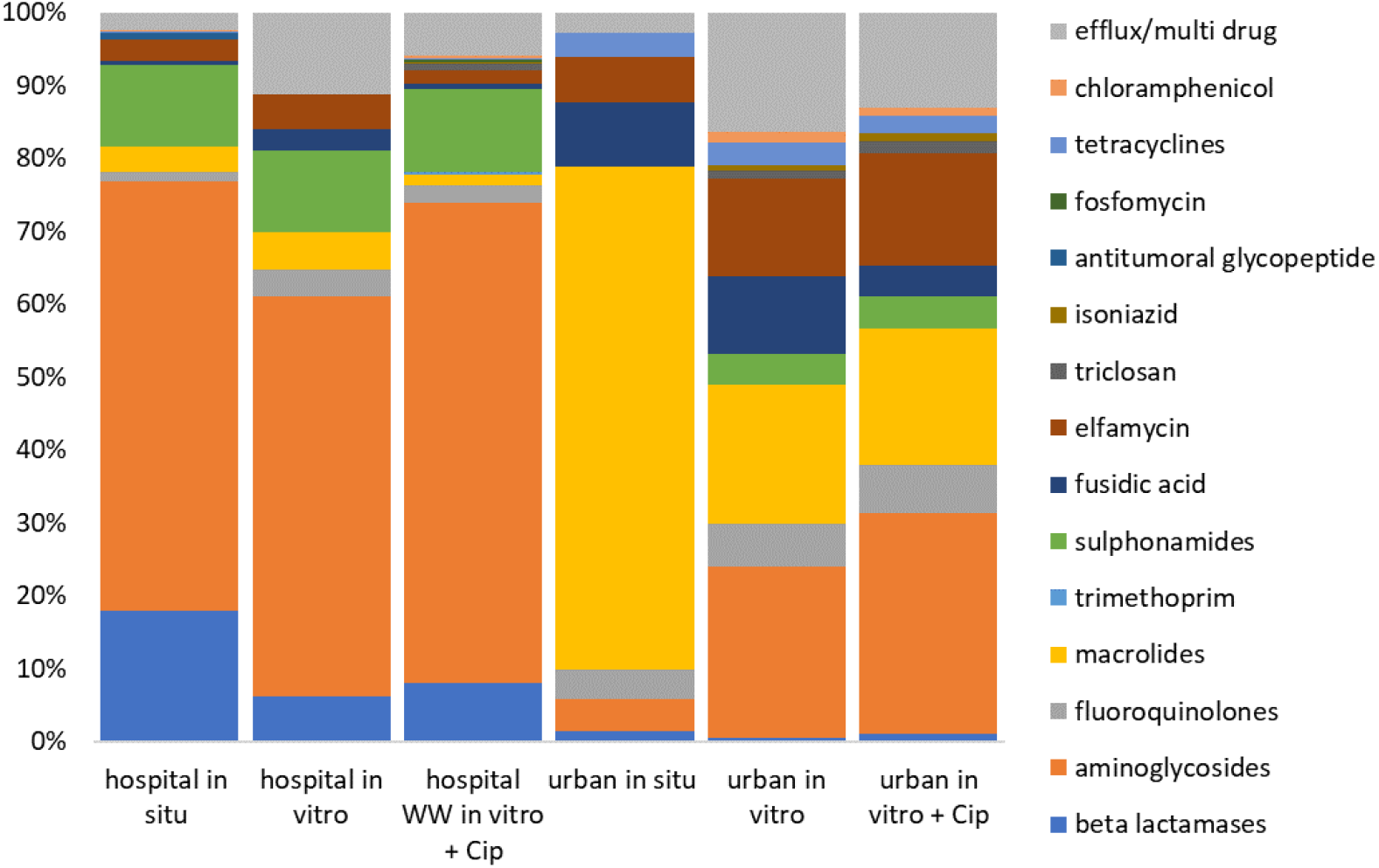
ARG metatranscriptome based on the CARD database. Mean numbers of mapped reads per antimicrobial resistance gene were summed by ARG class and are displayed per sample.

Overall the composition of the ARG metatranscriptome of the CARD resistome analysis correlated to the targeted resistome analysis (Supplementary Figure 5, Supplementary Table 4 and 9). Comparing the different environments for hospital biofilms (*in situ* vs *in vitro*), higher gene expression levels were detected for the *in situ* hospital biofilm between 3.2-fold and 15.2-fold (macrolides and beta-lactamases respectively). For the *in situ* vs *in vitro* urban biofilms, expression levels were lower for all detected ARG classes, apart from macrolide ARGs, which were expressed at a higher level in urban *in situ* biofilm (Supplementary Figure 9, Supplementary Table 4 and 9). For hospital *in situ* biofilm, six ARG classes had between 1.8 and 81.2-fold higher expression levels (genes encoding resistance to fluoroquinolones and aminoglycosides respectively) vs urban *in situ* biofilm. Comparing hospital *in vitro* biofilm exposed to ciprofloxacin to its control, overall, increasing fold-change for seven ARG classes, between 3- and 7.5-fold change for genes encoding for multi-drug efflux pumps and genes conferring resistance to beta-lactams, respectively, was detected. Interestingly, 5 ARG classes, namely aminoglycosides, beta lactams, quinolones, sulphonamides, and genes encoding for multi-drug efflux pumps, were detected to have higher gene expression levels in both, the targeted and non-targeted resistome approach, for the hospital *in vitro* biofilm exposed to ciprofloxacin compared to its control (Supplementary Figures 8 and 9, Supplementary Table 4, 8 and 9). For the urban *in vitro* biofilm exposed to ciprofloxacin, expression levels are lower, expect for genes encoding for beta-lactams were a slight increase (1.18-fold) could be detected compared to its control (Supplementary Figure 9, Supplementary Table 4 and 9).

### The active resistance mobilome

To study the expression of MGEs and more specifically MGEs associated with ARGs, assembled contigs were mapped against the plasmid sequence collection v0.4 available at the ACLAME database [44]. Subsequently, plasmid associated contigs identified by this approach were mapped back to the CARD database to identify specifically plasmids that carry ARGs. These ARG associated plasmids are referred to as the “active resistance mobilome” in the respective samples.

Global expression of plasmids was very low in all the samples studied (0.48% of all assembled contigs for hospital *in situ* biofilm and 0.14% for urban *in situ* biofilm (Supplementary Table 1)). In total, we counted respectively 115 and 66 plasmid-ARG associated contigs for the hospital and urban *in situ* biofilms and identified a diverse range of expressed plasmids (71 individual plasmids identified across all samples) expressing a diverse range of ARGs that could be grouped into 10 classes conferring resistance to antimicrobials, quaternary ammonium compounds and heavy metals. The highest expression level for the active resistance mobilome was observed in the *in vitro* hospital biofilm exposed to ciprofloxacin (Supplementary Figure 11).

55 different plasmids carrying a broad range of ARGs were identified as the active resistance mobilome across all hospital WW biofilm samples. These plasmids were mainly resistance plasmids found in Gram-negative pathogens such as the *bla-_VIM7_* carrying plasmid *pmATVIM-7* first identified in *P. aeruginosa* [50]; the IncP-6 plasmid *Rms149*, a small resistant plasmid associated with *P. aeruginosa* [58] (detected only in hospital *in situ* biofilm); several small resistance plasmids (e.g. *pKMA757, pARD3079, pKMA202*, and *pKMA5*) associated with *Actinobacilli* [59] and plasmid families representing the groups IncP, F, N and Q (Supplementary Figure 11). They primarily expressed aminoglycoside, sulphonamide, chloramphenicol, and beta-lactam resistance genes (Supplementary Figure 11, Figure 6). 14 plasmids were detected in all hospital WW biofilm samples, and 7 additional plasmids were only shared between the *in situ* hospital WW biofilms and the *in vitro* hospital WW biofilm exposed to ciprofloxacin (Supplementary Figure 11). Interestingly, 22 plasmids, carrying between 1 and 5 ARGs, were only identified in the *in vitro* hospital WW biofilm exposed to ciprofloxacin (Supplementary Figure 11).

**Figure 6:**
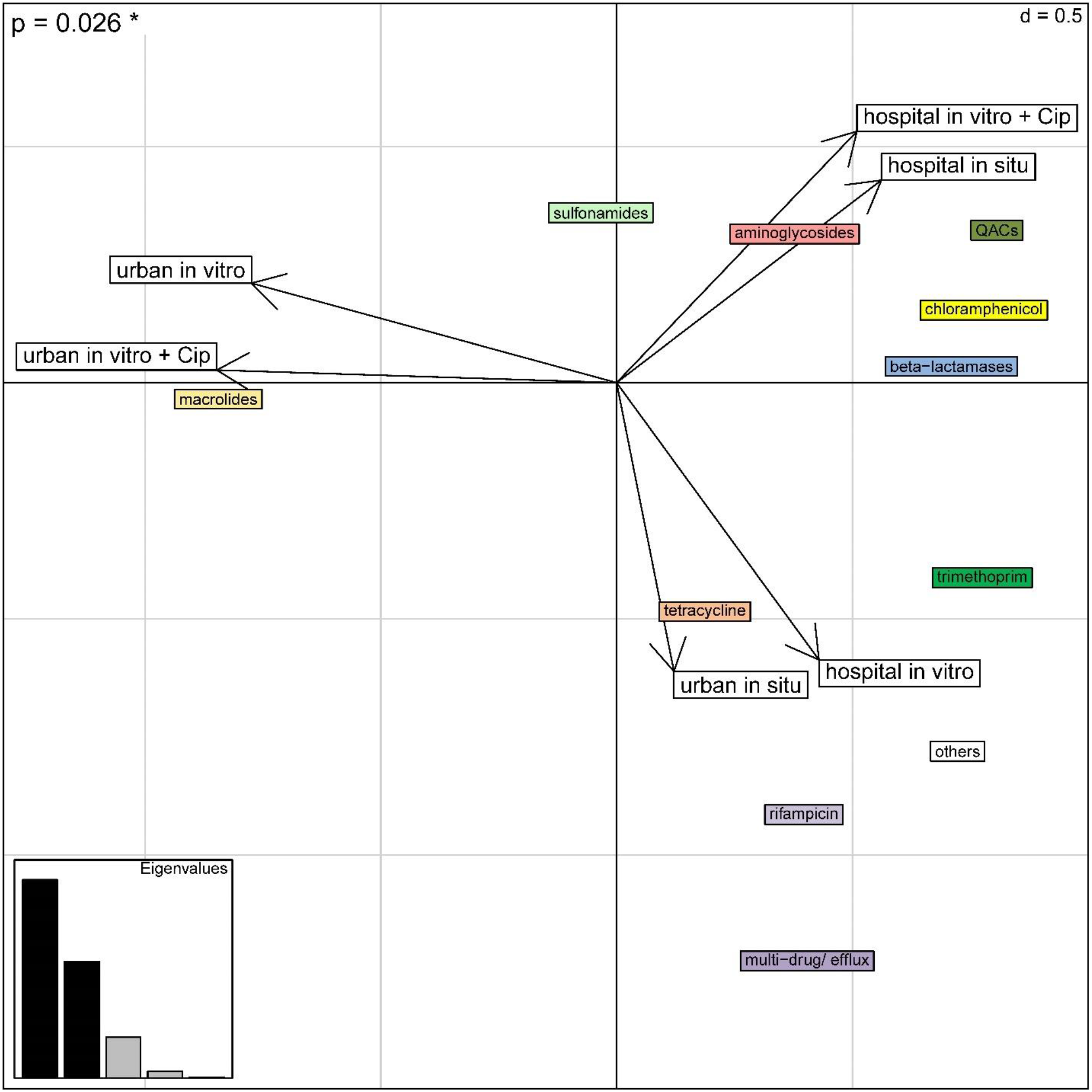
Between correspondence analysis (bCA) assessing the association between the type of ARGs classes present on plasmids, in terms of ARG class abundance (numbers of contigs) and the biofilm origin.

14 plasmids were unique for urban WW biofilm samples (*in situ* and *in vitro*). These plasmids mainly carried macrolide and tetracycline resistance genes found in Gram-positive bacteria, such as Lactobacilli (56). In addition, the *Enterococcus faecium* and *faecalis* plasmids *pRUM* and *pTEF1* were identified as expressing both the macrolide resistance gene *ermB* [60–62] (Supplementary Figure 11). The broad host range IncP-1 plasmid pKJK5 was also detected, expressing an aminoglycoside (*aadA6*) and a sulphonamide (*sul1*) resistance gene [63,64]. One Gram-negative multidrug resistance plasmid *pJR1* [65] was detected, carrying the chloramphenicol resistance gene *catB2* and the tetracycline resistance gene *tet(G*) (Supplementary Figure 11). Fifteen plasmids were uniquely identified in the *in situ* urban WW biofilm and not in its *in vitro* counterpart. Five of these plasmids were also detected in the hospital WW biofilms expressing different ARGs (e.g. *pJHCMW1* and *R46*), whereas they only carried a single ARG when detected in the *in situ* urban biofilm (Supplementary Figure 11).

In the hospital WW biofilms, many plasmids were identified to carry multiple resistance genes (up to 5 ARGs carried on one MGE contig, Supplementary Figure 11). These genes mainly encoded resistance to aminoglycosides, beta-lactams, sulphonamides, chloramphenicol, and quaternary ammonium compounds (Figure 6, Supplementary Figure 11). For the urban WW biofilms, MGEs that express multiple resistance genes were also identified, but with less ARGs associated (maximum three ARGs on one plasmid in the *in situ* urban WW biofilm) (Supplementary Figure 11). The ARGs mainly detected in the active resistance mobilome of urban WW biofilms were macrolide, tetracyclines and multi-drug efflux pumps (Figure 6, Supplementary Figure 11). Contrary to what is observed for the effect of ciprofloxacin on the active resistance mobilome of the *in vitro* hospital biofilm, less MGE-ARG associated contigs were identified, and a much lower diversity of ARGs were expressed in the *in vitro* urban biofilm exposed to ciprofloxacin (mainly macrolide resistance genes) (Supplementary Figure 11, Figure 6).

Overall, correspondence analysis (CA) of our mobilome data, revealed a prominent association between the ARGs present on plasmids and their biofilm origin (Cramér’s V statistic = 0.604; χ2-test *p*-value < 0.001; Supplementary Figure 12). To assess if this association was also observed at the level of the ARGs classes, we applied a between CA (bCA). This recovered 33% of the total association between biofilm origin and the ARGs composition of the mobilome (*p*-value=0.026; Figure 6).

## Discussion

The goal of this study was to develop a comprehensive experimental approach to assess WW biofilms as putative hotspots for actual transfer of ARGs via MGEs within these communities using metagenomic approaches. We furthermore wanted to quantify and visualize the EPS components of these biofilms, due to the proposed role of the EPS in horizontal gene transfer within biofilms, as well as their relevance for biofilms to tolerate antibiotics and exogeneous stressors [29,31,66–68].

Here we introduced a method by confocal laser scanning microscopy (CLSM) to visualize and quantify the EPS of hospital and urban WW biofilms allowing to study EPS components in these biofilms over time and under different anthropogenic constraints. We showed that i) the EPS components investigated are differentially abundant dependent on their growth environment (*in* situ and *in vitro*) and their sampling origin (hospital vs urban WW) and ii) ciprofloxacin at MSC impacted the quantity of the measured EPS components. The decrease in the quantity of polysaccharides in the urban and in hospital *in vitro* biofilms, and for eDNA in the hospital *in vitro* biofilm, due to the presence of ciprofloxacin might be linked to a regulatory response of the biofilm community. Indeed, it has been shown that exogeneous stress in multi-species biofilms leads to decreased EPS production [69]. *In vitro* eDNA was proportionally more abundant in hospital WW biofilms compared to urban WW biofilms, which could be linked to higher amounts of surfactants, antibiotics, and other pharmaceutical and chemical pollution in hospital WW [7] that in turn may enhance bacterial lysis and active eDNA secretion in those biofilms [70]. eDNA is known to promote tolerance to antimicrobial agents in biofilms by inhibiting the diffusion of cationic molecules [71] and by increasing the resistance to aminoglycosides in *Pseudomonas aeruginosa* biofilms [72,73]. It has also been shown that eDNA is involved in transfer of the ARG *tetM* by the conjugative transposon Tn916 [68], through transformation. Transformation is suspected to be an important mechanism for gene transfer between bacterial communities in biofilms [68]. We showed that the microbiota of hospital and urban WW biofilms is distinct, which is likely correlated to the different organization and relative abundances detected for the EPS in the two WW biofilm niches, as shown by our analysis (Supplementary Figure 5 and Supplementary Table 6). The significant increase of dead cells observed for the *in vitro* hospital WW biofilm exposed to ciprofloxacin could be linked to the predominant Gram-negative genera in hospital WW biofilms (Figure 3), which are highly susceptible to ciprofloxacin.

By implementing a meta-transcriptomic approach, at first, we wanted to assess the basal differential expression levels of mobile genetic elements (MGEs) associated with ARGs as a proxy for actual transfer of ARGs within hospital and urban WW biofilms. Second, we wanted to compare these levels to *in vitro* reconstituted biofilms and third, we wanted to see whether these “basal” expression levels do change in response to a single antibiotic added to the WW environment. We showed that a) basal ARG and MGEs expression levels are higher in *in situ* than *in vitro* WW hospital biofilms; b) expression levels of ARGs and MGEs increase upon adding a minimal selective concentration (MSC) of ciprofloxacin to the *in vitro* hospital WW biofilm environment and c) that expression of plasmids carrying specific ARG classes and multiple ARGs at the same time, is associated with the biofilm origin (Figure 6). We were able to detect a broad range of plasmids that were uniquely detected in either, hospital or urban WW biofilms. Interestingly, plasmids detected in both, hospital and urban *in situ* and *in vitro* biofilms, carried different types and quantities of ARGs (Supplementary Figure 9), suggesting that the plasmids cargo (content of ARGs) is adapted to its environment. Hence, we showed that the nature of ARGs present on plasmids is correlated to the biofilm environment and origin.

Interestingly, the reconstituted *in vitro* hospital and urban WW biofilms were more like each other concerning their microbiota composition than their natural (*in situ*) biofilm counterparts, which was not the case for the resistome, resistome expression nor active resistance mobilome. This surprising result suggests that the *in vitro* laboratory conditions have a larger impact on the biofilm community structure than the WW in which they developed. This questions the ability of our experimental set-up to implement microcosms in the laboratory to mimic the natural environment. Further optimization of the *in vitro* model is needed, which could be achieved for example by more frequent refreshment of the WW alimentation in the *in vitro* system, or better control of the oxygen levels.

Among the classifiable rRNA reads from the metatranscriptomic analysis we identified a large majority of *Enterobacteriaceae* in the hospital biofilms, and a mix of *Flavobacteriaceae, Enterobacteriaceae, Streptococaceae* and *Francisellaceae* in the urban *in situ* biofilm (Supplementary Figure 6 and Supplementary Table 2). The fact that these rRNA reads were obtained from the RNA sequencing run may indicate that these families were highly “active” in these biofilms although not highly abundant based on 16S rDNA sequencing (Figure 3). The highly active proportion of *Enterobacteriaceae* in hospital *in situ* and *in vitro* biofilms might indeed explain the higher expression levels and the types of MGEs and ARGs detected in these samples, and their response to antibiotic stress. This further highlights the need to study the ‘active biofilm community’ and might point towards the fact that the key players for accumulation and transfer of ARGs and MGEs in highly anthropized sites such as WW are indeed human-gut associated bacteria.

In a recent study, the Plascad tool was designed for plasmid classification, AMR gene annotation, and plasmid visualization [74] and they showed that: i) most plasmid-borne ARGs, including those localized on class-1 integrons, are enriched in conjugative plasmids in their datasets, and ii) transfer between AMR plasmids and bacterial chromosomes was mediated by insertion sequences, mainly those belonging to the IS6 family, IS26 and IS6100 [74]. We have shown previously that IS26 and IS6100 were both abundant in WW [7] and in this study we show that IS6100 is highly expressed in hospital WW biofilms. IS26 is frequently detected in plasmids of clinical importance associated with antibiotic resistance genes. Furthermore, IS26 has also been associated with the expression of antibiotic resistance genes [75]. Finally, the distinct plasmid profiles (Figure 6 and Supplementary Figure 11) and their specific ARG expression profiles for hospital and urban WW biofilms point towards a strong environmental selection pressure shaping the active resistance mobilome.

Here we provide an approach to comprehensively study multi-species biofilms that form in naturally as well as highly anthropized environments such as WW. We show that hospital and urban wastewaters shape the structure and active resistome of environmental biofilms, and we confirmed that hospital WW is an important hot spot for the dissemination and selection of AMR. Hence, we propose that studying hospital WW biofilms may present an ideal model to further unravel the complex correlation between pollution and AMR selection and the structural, ecological, functional, and genetic organization of multi-species biofilms as antibiotic resistance factories. Finally, our results highlight that for all studied biofilm samples, resistome and mobilome significantly depended on their environment (*in vitro* vs *in situ*) and origin (WW).

## Ethics approval and Consent to participate

Not applicable.

## Consent for publication

All authors gave their consent for publication.

## Availability of data and materials

16S rRNA sequence data are available at the European Nucleotide Archive (ENA) under the accession numbers ERS14475391-8. RNA sequencing data (metatranscriptomic data) will be available at the NCBI under project number PRJNA930611. All other important raw data needed to reconstruct the findings of our study are made available in the supplementary material.

## Authors’ contributions

C.D., S. D.-R., M-C.P., and E.B. designed the study, Ca. D. and S.A. developed the metatranscriptomic analysis pipeline, M.G. and E.B. performed experiments, E.B., Ca.D., H.M.H., T.J. and S.P.K. performed data analysis, O.C. and H.M.H provided critical comments on the content of the manuscript, E.B., wrote the manuscript with contribution of all other co-authors. All authors reviewed the manuscript.

## Acknowledgements

Not applicable.

## References

1. Balcazar JL, Subirats J, Borrego CM. The role of biofilms as environmental reservoirs of antibiotic resistance. FrontMicrobiol. 2015;6:1216.

2. Hennequin C, Aumeran C, Robin F, Traore O, Forestier C. Antibiotic resistance and plasmid transfer capacity in biofilm formed with a CTX-M-15-producing *Klebsiella pneumoniae* isolate. J Antimicrob Chemother. 2012;67:2123–30.

3. Ferreira C, Veldhoen M. Host and microbes date exclusively. Cell. 2012;149:1428–30.

4. Abe K, Nomura N, Suzuki S. Biofilms: hot spots of horizontal gene transfer (HGT) in aquatic environments, with a focus on a new HGT mechanism. FEMS Microbiology Ecology. 2020;96. Available from: https://doi.org/10.1093/femsec/fiaa031

5. Allen HK, Donato J, Wang HH, Cloud-Hansen KA, Davies J, Handelsman J. Call of the wild: antibiotic resistance genes in natural environments. NatRevMicrobiol. 2010;8:251–9.

6. Flandroy L, Poutahidis T, Berg G, Clarke G, Dao MC, Decaestecker E, et al. The impact of human activities and lifestyles on the interlinked microbiota and health of humans and of ecosystems. SciTotal Environ. 2018;627:1018–38.

7. Buelow E, Rico A, Gaschet M, Lourenço J, Kennedy SP, Wiest L, et al. Hospital discharges in urban sanitation systems: Long-term monitoring of wastewater resistome and microbiota in relationship to their eco-exposome. Water Research X. 2020;7:100045.

8. Harris S, Morris C, Morris D, Cormican M, Cummins E. The effect of hospital effluent on antimicrobial resistant *E. coli* within a municipal wastewater system. EnvironSciProcessImpacts. 2013;15:617–22.

9. Jia A, Wan Y, Xiao Y, Hu J. Occurrence and fate of quinolone and fluoroquinolone antibiotics in a municipal sewage treatment plant. Water Research. 2012;46:387–94.

10. Hayes A, May Murray L, Catherine Stanton I, Zhang L, Snape J, Hugo Gaze W, et al. Predicting selection for antimicrobial resistance in UK wastewater and aquatic environments: Ciprofloxacin poses a significant risk. Environment International. 2022;169:107488.

11. Kim D, Nguyen LN, Oh S. Ecological impact of the antibiotic ciprofloxacin on microbial community of aerobic activated sludge. Environ Geochem Health. 2020;42:1531–41.

12. Nakata H, Kannan K, Jones PD, Giesy JP. Determination of fluoroquinolone antibiotics in wastewater effluents by liquid chromatography–mass spectrometry and fluorescence detection. Chemosphere. 2005;58:759–66.

13. Beaber JW, Hochhut B, Waldor MK. SOS response promotes horizontal dissemination of antibiotic resistance genes. Nature. Nature Publishing Group; 2004;427:72–4.

14. Guerin É, Cambray G, Sanchez-Alberola N, Campoy S, Erill I, Re SD, et al. The SOS Response Controls Integron Recombination. Science. American Association for the Advancement of Science; 2009;324:1034–1034.

15. Hu Y, Yang X, Qin J, Lu N, Cheng G, Wu N, et al. Metagenome-wide analysis of antibiotic resistance genes in a large cohort of human gut microbiota. NatCommun. 2013;4:2151.

16. Forslund K, Sunagawa S, Coelho LP, Bork P. Metagenomic insights into the human gut resistome and the forces that shape it. Bioessays. 2014;36:316–29.

17. Sommer MO, Church GM, Dantas G. The human microbiome harbors a diverse reservoir of antibiotic resistance genes. Virulence. 2010;1:299–303.

18. Karkman A, Parnanen K, Larsson DGJ. Fecal pollution can explain antibiotic resistance gene abundances in anthropogenically impacted environments. NatCommun. 2019;10:80-018-07992–3.

19. Pehrsson EC, Forsberg KJ, Gibson MK, Ahmadi S, Dantas G. Novel resistance functions uncovered using functional metagenomic investigations of resistance reservoirs. FrontMicrobiol. 2013;4:145.

20. Singer AC, Shaw H, Rhodes V, Hart A. Review of Antimicrobial Resistance in the Environment and Its Relevance to Environmental Regulators. FrontMicrobiol. 2016;7:1728.

21. McInnes RS, uz-Zaman MH, Alam IT, Ho SFS, Moran RA, Clemens JD, et al. Metagenome-Wide Analysis of Rural and Urban Surface Waters and Sediments in Bangladesh Identifies Human Waste as a Driver of Antibiotic Resistance. mSystems. American Society for Microbiology; 2021;6:e00137–21.

22. Manaia CM, Rocha J, Scaccia N, Marano R, Radu E, Biancullo F, et al. Antibiotic resistance in wastewater treatment plants: Tackling the black box. EnvironInt. 2018;115:312–24.

23. Rizzo L, Manaia C, Merlin C, Schwartz T, Dagot C, Ploy MC, et al. Urban wastewater treatment plants as hotspots for antibiotic resistant bacteria and genes spread into the environment: a review. SciTotal Environ. 2013;447:345–60.

24. Haenni M, Dagot C, Chesneau O, Bibbal D, Labanowski J, Vialette M, et al. Environmental contamination in a high-income country (France) by antibiotics, antibiotic-resistant bacteria, and antibiotic resistance genes: Status and possible causes. Environ Int. 2022;159:107047.

25. Luo Y-H, Lai YS, Zheng C, Ilhan ZE, Ontiveros-Valencia A, Long X, et al. Increased expression of antibiotic-resistance genes in biofilm communities upon exposure to cetyltrimethylammonium bromide (CTAB) and other stress conditions. Science of The Total Environment. 2021;765:144264.

26. Proia L, von Schiller D, Sànchez-Melsió A, Sabater S, Borrego CM, Rodríguez-Mozaz S, et al. Occurrence and persistence of antibiotic resistance genes in river biofilms after wastewater inputs in small rivers. Environmental Pollution. 2016;210:121–8.

27. Flemming H-C, Wuertz S. Bacteria and archaea on Earth and their abundance in biofilms. Nat Rev Microbiol. 2019;17:247–60.

28. Ratajczak M, Kaminska D, Dlugaszewska J, Gajecka M. Antibiotic Resistance, Biofilm Formation, and Presence of Genes Encoding Virulence Factors in Strains Isolated from the Pharmaceutical Production Environment. Pathogens. Multidisciplinary Digital Publishing Institute; 2021;10:130.

29. Flemming HC, Wingender J. The biofilm matrix. NatRevMicrobiol. 2010;8:623–33.

30. Xiao J, Koo H. Structural organization and dynamics of exopolysaccharide matrix and microcolonies formation by Streptococcus mutans in biofilms. JApplMicrobiol. 2010;108:2103–13.

31. Soares A, Roussel V, Pestel-Caron M, Barreau M, Caron F, Bouffartigues E, et al. Understanding Ciprofloxacin Failure in *Pseudomonas aeruginosa* Biofilm: Persister Cells Survive Matrix Disruption. Front Microbiol. 2019;10:2603.

32. Buelow E, Bayjanov JR, Majoor E, Willems RJL, Bonten MJM, Schmitt H, et al. Limited influence of hospital wastewater on the microbiome and resistome of wastewater in a community sewerage system. FEMS MicrobiolEcol. 2018;

33. Chonova T, Lecomte V, Bertrand-Krajewski JL, Bouchez A, Labanowski J, Dagot C, et al. The SIPIBEL project: treatment of hospital and urban wastewater in a conventional urban wastewater treatment plant. EnvironSciPollutResInt. 2018;25:9197–206.

34. Bengtsson-Palme J, Larsson DG. Concentrations of antibiotics predicted to select for resistant bacteria: Proposed limits for environmental regulation. EnvironInt. 2016;86:140–9.

35. Buelow E, Bello Gonzalez TDJ, Fuentes S, de Steenhuijsen Piters WAA, Lahti L, Bayjanov JR, et al. Comparative gut microbiota and resistome profiling of intensive care patients receiving selective digestive tract decontamination and healthy subjects. Microbiome. 2017;5:88-017-0309-z.

36. Langsrud S, Sundheim G, Holck AL. Cross-resistance to antibiotics of Escherichia coli adapted to benzalkonium chloride or exposed to stress-inducers. JApplMicrobiol. 2004;96:201–8.

37. Buffet-Bataillon S, Tattevin P, Bonnaure-Mallet M, Jolivet-Gougeon A. Emergence of resistance to antibacterial agents: the role of quaternary ammonium compounds--a critical review. IntJAntimicrobAgents. 2012;39:381–9.

38. Gillings MR, Gaze WH, Pruden A, Smalla K, Tiedje JM, Zhu YG. Using the class 1 integron-integrase gene as a proxy for anthropogenic pollution. ISME J. 2015;9:1269–79.

39. Wood DE, Salzberg SL. Kraken: ultrafast metagenomic sequence classification using exact alignments. Genome Biology. 2014;15:R46.

40. Pruesse E, Quast C, Knittel K, Fuchs BM, Ludwig W, Peplies J, et al. SILVA: a comprehensive online resource for quality checked and aligned ribosomal RNA sequence data compatible with ARB. Nucleic Acids Res. 2007;35:7188–96.

41. Solden L, Lloyd K, Wrighton K. The bright side of microbial dark matter: lessons learned from the uncultivated majority. Curr Opin Microbiol. 2016;31:217–26.

42. Evaluation of metatranscriptomic protocols and application to the study of freshwater microbial communities - Tsementzi - 2014 - Environmental Microbiology Reports: /doi/abs/10.1111/1758-2229.12180

43. Alcock BP, Raphenya AR, Lau TTY, Tsang KK, Bouchard M, Edalatmand A, et al. CARD 2020: antibiotic resistome surveillance with the comprehensive antibiotic resistance database. Nucleic Acids Res. 2020;48:D517–25.

44. Leplae R, Lima-Mendez G, Toussaint A. ACLAME: a CLAssification of Mobile genetic Elements, update 2010. Nucleic Acids Res. 2010;38:D57–61.

45. Liu Z, Klümper U, Liu Y, Yang Y, Wei Q, Lin J-G, et al. Metagenomic and metatranscriptomic analyses reveal activity and hosts of antibiotic resistance genes in activated sludge. Environ Int. 2019;129:208–20.

46. Okshevsky M, Meyer RL. Evaluation of fluorescent stains for visualizing extracellular DNA in biofilms. Journal of Microbiological Methods. 2014;105:102–4.

47. Hothorn T, Bretz F, Westfall P. Simultaneous inference in general parametric models. Biom J. 2008;50:346–63.

48. Harrison XA. Using observation-level random effects to model overdispersion in count data in ecology and evolution. PeerJ. PeerJ Inc.; 2014;2:e616.

49. Jakobsson M, Edge MD, Rosenberg NA. The Relationship Between FST and the Frequency of the Most Frequent Allele. Genetics. 2013;193:515–28.

50. Toleman MA, Rolston K, Jones RN, Walsh TR. blaVIM-7, an Evolutionarily Distinct Metallo-β-Lactamase Gene in a *Pseudomonas aeruginosa* Isolate from the United States. ANTIMICROB AGENTS CHEMOTHER. :48:329–332.

51. Manchanda V, Sanchaita S, Singh N. Multidrug Resistant Acinetobacter. J Glob Infect Dis. 2010;2:291–304.

52. Vandewalle JL, Goetz GW, Huse SM, Morrison HG, Sogin ML, Hoffmann RG, et al. *Acinetobacter, Aeromonas* and *Trichococcus* populations dominate the microbial community within urban sewer infrastructure. EnvironMicrobiol. 2012;14:2538–52.

53. Fisher JC, Levican A, Figueras MJ, McLellan SL. Population dynamics and ecology of *Arcobacter* in sewage. FrontMicrobiol. 2014;5:525.

54. Ferreira S, Fraqueza MJ, Queiroz JA, Domingues FC, Oleastro M. Genetic diversity, antibiotic resistance and biofilm-forming ability of *Arcobacter butzleri* isolated from poultry and environment from a Portuguese slaughterhouse. IntJFood Microbiol. 2013;162:82–8.

55. Batovska J, Mee PT, Lynch SE, Sawbridge TI, Rodoni BC. Sensitivity and specificity of metatranscriptomics as an *arbovirus* surveillance tool. Sci Rep. Nature Publishing Group; 2019;9:19398.

56. Sivalingam P, Poté J, Prabakar K. Extracellular DNA (eDNA): Neglected and Potential Sources of Antibiotic Resistant Genes (ARGs) in the Aquatic Environments. Pathogens. 2020;9:874.

57. Eramo A, Medina WRM, Fahrenfeld NL. Viability-based quantification of antibiotic resistance genes and human fecal markers in wastewater effluent and receiving waters. The Science of the total environment. NIH Public Access; 2019;656:495.

58. Haines AS, Jones K, Cheung M, Thomas CM. The *IncP-6* plasmid *Rms149* consists of a small mobilizable backbone with multiple large insertions. J Bacteriol. 2005;187:4728–38.

59. Matter D, Rossano A, Sieber S, Perreten V. Small multidrug resistance plasmids in *Actinobacillus porcitonsillarum*. Plasmid. 2008;59:144–52.

60. Jensen LB, Frimodt-Moller N, Aarestrup FM. Presence of erm gene classes in gram-positive bacteria of animal and human origin in Denmark. FEMS MicrobiolLett. 1999;170:151–8.

61. Jensen LB, Garcia-Migura L, Valenzuela AJS, Løhr M, Hasman H, Aarestrup FM. A classification system for plasmids from *enterococci* and other Gram-positive bacteria. Journal of Microbiological Methods. 2010;80:25–43.

62. Palmer KL, Kos VN, Gilmore MS. Horizontal Gene Transfer and the Genomics of *Enterococcal* Antibiotic Resistance. Curr Opin Microbiol. 2010;13:632–9.

63. Popowska M, Krawczyk-Balska A. Broad-host-range *IncP-1* plasmids and their resistance potential. Frontiers in Microbiology. 2013;4;Article 44.

64. Røder HL, Hansen LH, Sørensen SJ, Burmølle M. The impact of the conjugative *IncP-1* plasmid *pKJK5* on multispecies biofilm formation is dependent on the plasmid host. FEMS Microbiol Lett. 2013;344:186–92.

65. Wu J-R, Shieh HK, Shien J-H, Gong S-R, Chang P-C. Molecular characterization of plasmids with antimicrobial resistant genes in avian isolates of *Pasteurella multocida*. Avian Dis. 2003;47:1384–92.

66. Karygianni L, Ren Z, Koo H, Thurnheer T. Biofilm Matrixome: Extracellular Components in Structured Microbial Communities. Trends in Microbiology. 2020;28:668–81.

67. Gu C, Gao P, Yang F, An D, Munir M, Jia H, et al. Characterization of extracellular polymeric substances in biofilms under long-term exposure to ciprofloxacin antibiotic using fluorescence excitation-emission matrix and parallel factor analysis. Environ Sci Pollut Res. 2017;24:13536–45.

68. Hannan S, Ready D, Jasni AS, Rogers M, Pratten J, Roberts AP. Transfer of antibiotic resistance by transformation with eDNA within oral biofilms. FEMS Immunology & Medical Microbiology. 2010;59:345–9.

69. Loustau E, Leflaive J, Boscus C, Amalric Q, Ferriol J, Oleinikova O, et al. The Response of Extracellular Polymeric Substances Production by Phototrophic Biofilms to a Sequential Disturbance Strongly Depends on Environmental Conditions. Frontiers in Microbiology. 2021;12; article 742027.

70. Okshevsky M, Meyer RL. The role of extracellular DNA in the establishment, maintenance and perpetuation of bacterial biofilms. Critical Reviews in Microbiology. Taylor & Francis; 2015;41:341–52.

71. Wilton M, Charron-Mazenod L, Moore R, Lewenza S. Extracellular DNA Acidifies Biofilms and Induces Aminoglycoside Resistance in *Pseudomonas aeruginosa*. Antimicrobial Agents and Chemotherapy. American Society for Microbiology; 2015;60:544–53.

72. Doroshenko N, Tseng BS, Howlin RP, Deacon J, Wharton JA, Thurner PJ, et al. Extracellular DNA impedes the transport of vancomycin in *Staphylococcus epidermidis* biofilms preexposed to subinhibitory concentrations of vancomycin. AntimicrobAgents Chemother. 2014;58:7273–82.

73. Lewenza S. Extracellular DNA-induced antimicrobial peptide resistance mechanisms in *Pseudomonas aeruginosa*. Front Microbiol. 2013;4:21.

74. Che Y, Yang Y, Xu X, Břinda K, Polz MF, Hanage WP, et al. Conjugative plasmids interact with insertion sequences to shape the horizontal transfer of antimicrobial resistance genes. Proc Natl Acad Sci USA. 2021;118:e2008731118.

75. Varani A, He S, Siguier P, Ross K, Chandler M. The *IS6* family, a clinically important group of insertion sequences including *IS26*. Mobile DNA. 2021;12:11.

