## Supplementary Figures 1-10 for "Hospital and urban wastewaters shape the structure and active resistome of environmental biofilms"

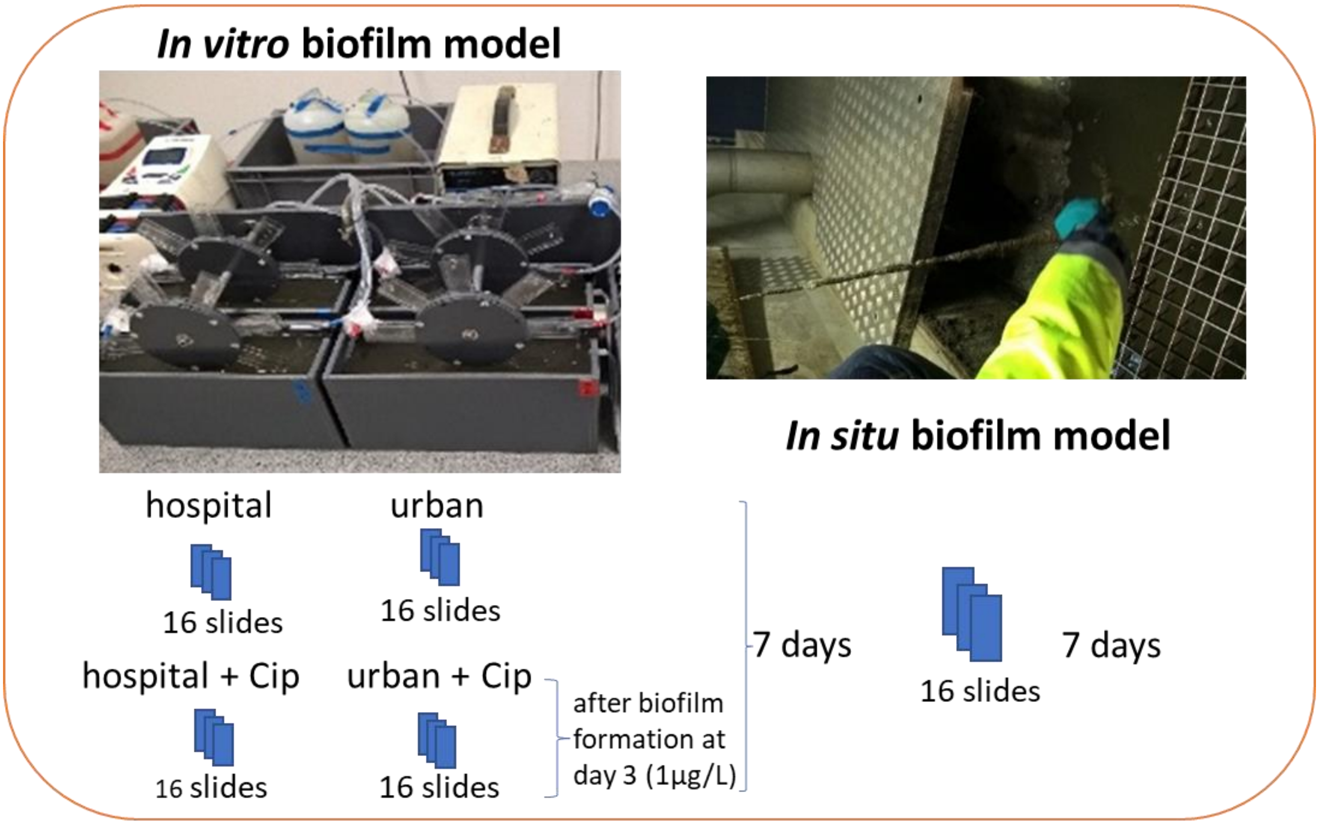


**Supplementary Figure 1:** Biofilm *in vitro* and *in situ* model.

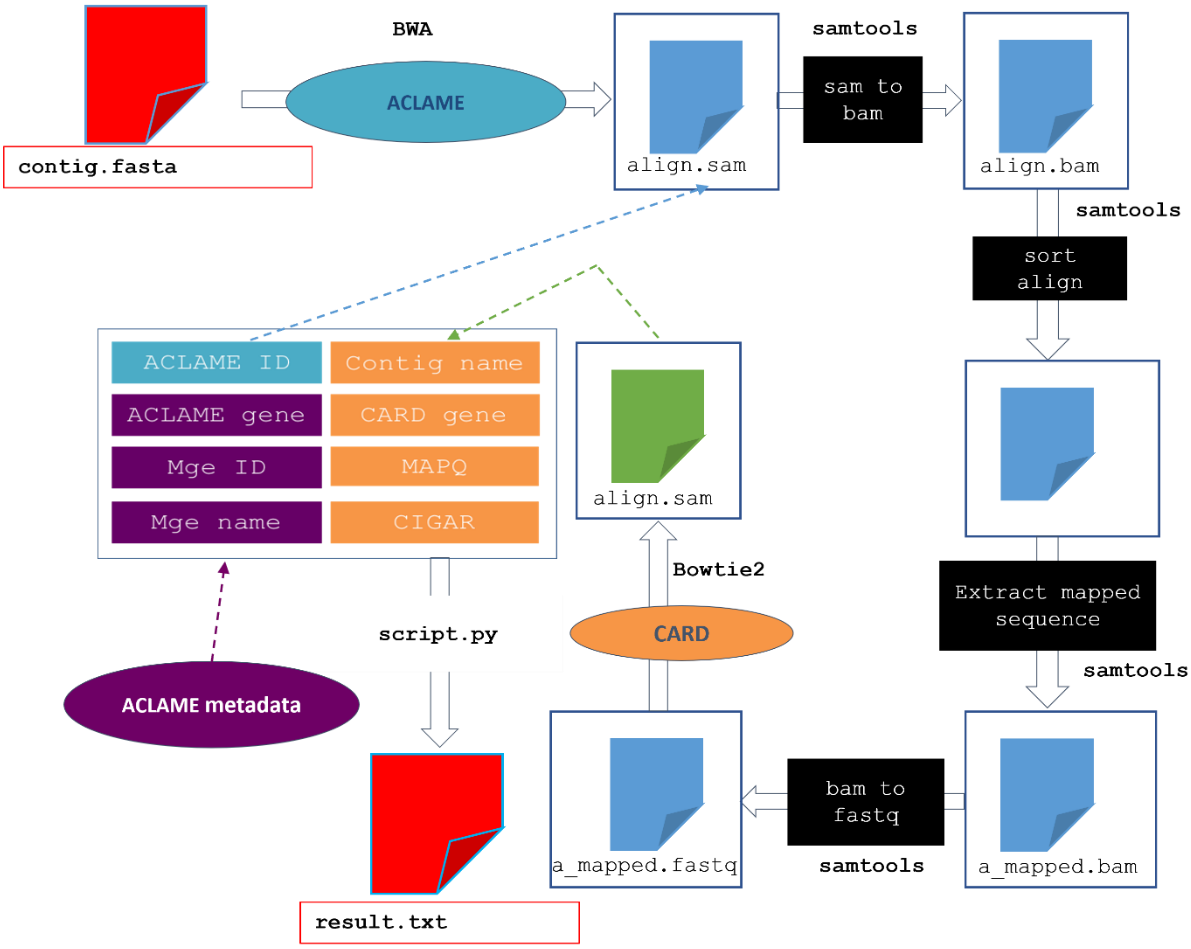


**Supplementary Figure 2: The ARoM pipeline.** ARoM (ARg On Mobilome) performs requests on the ACLAME database to retrieve MGEs and the CARD database to retrieve ARGs. Applied on data extracted from ACLAME, the requests on CARD allowed to detect ARG-carrying MGEs. The final step of ARoM merged data and metadata from CARD and ACLAME and their matched values. The outfile obtained allowed expert analysis of the results.

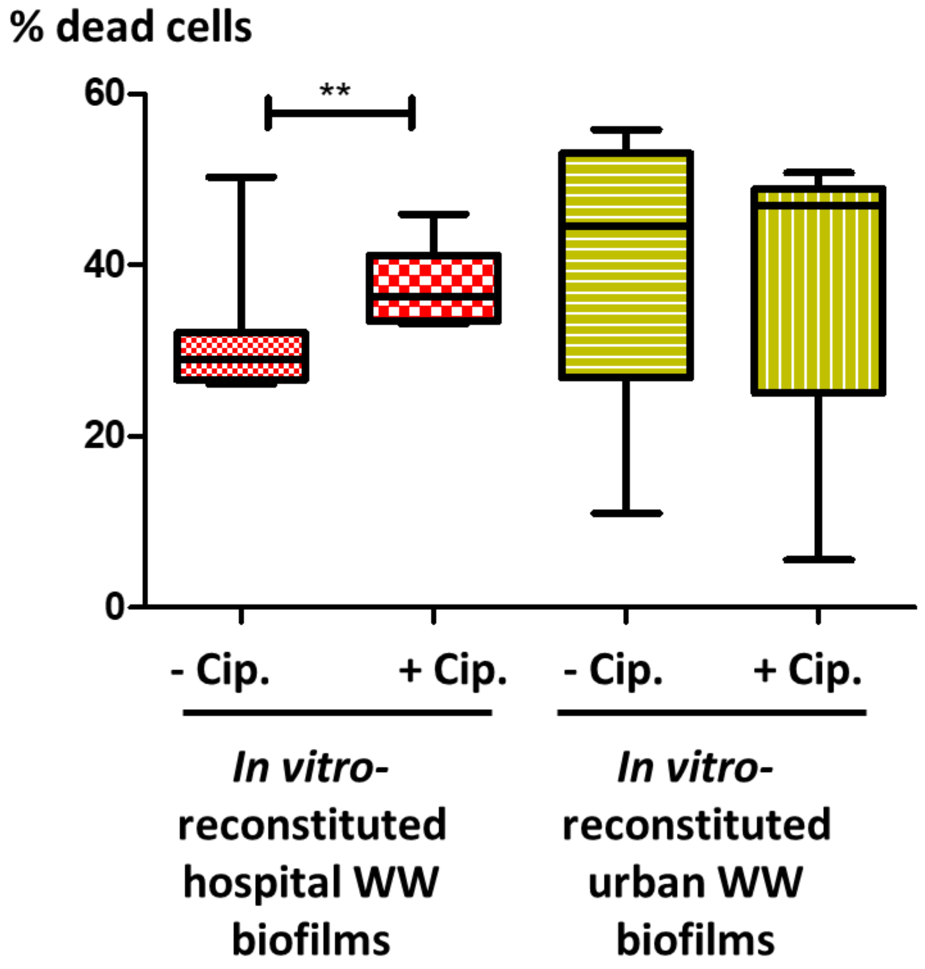


**Supplementary Figure 3**: Percentage of dead cells in *in vitro* biofilms grown in hospital and urban wastewaters. +Ab = ciprofloxacin was added at 1mg/L at day3, daily until biofilm collection and live-dead staining at day 7.

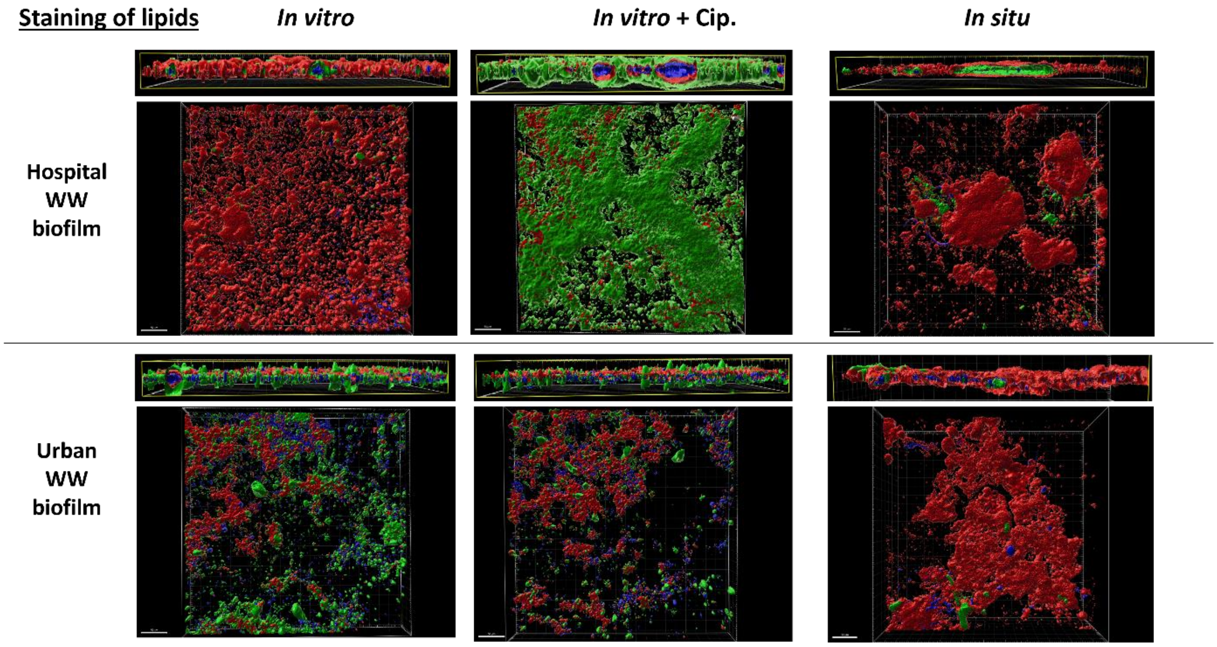


**Supplementary Figure 4a:** 3D imaging of lipids (red) and proteins (green), for hospital and urban *in-situ* and *in-vitro* biofilms. Autofluorescence for unknown compounds is in blue. +Ab= *in vitro* biofilms were exposed after 3 days of initial biofilm formation with 1µg/ml ciprofloxacin for 4 consecutive days and harvested after exposure of 4 hours on the final day.

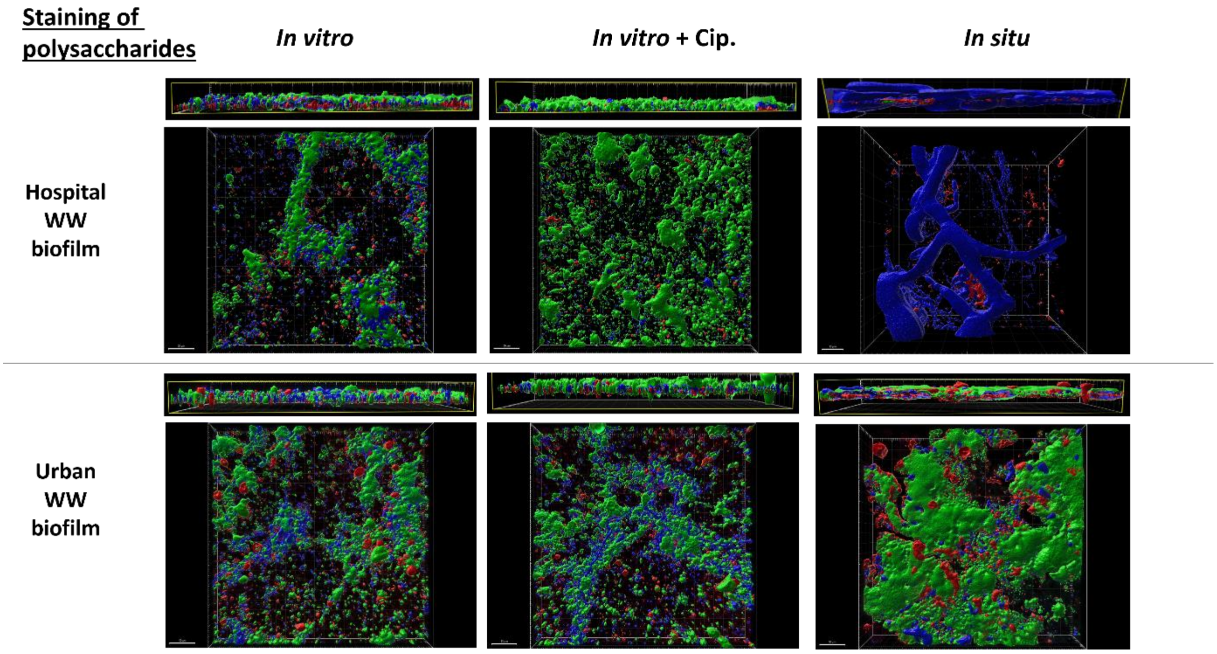


**Supplementary Figure 4b:** 3D imaging of polysaccharides (red) and proteins (green), for hospital and urban *in-situ* and *in-vitro* biofilms. Autofluorescence for unknown compounds is in blue. +Ab= *in vitro* biofilms were exposed after 3 days of initial biofilm formation with 1µg/ml ciprofloxacin for 4 consecutive days and harvested after exposure of 4 hours on the final day.

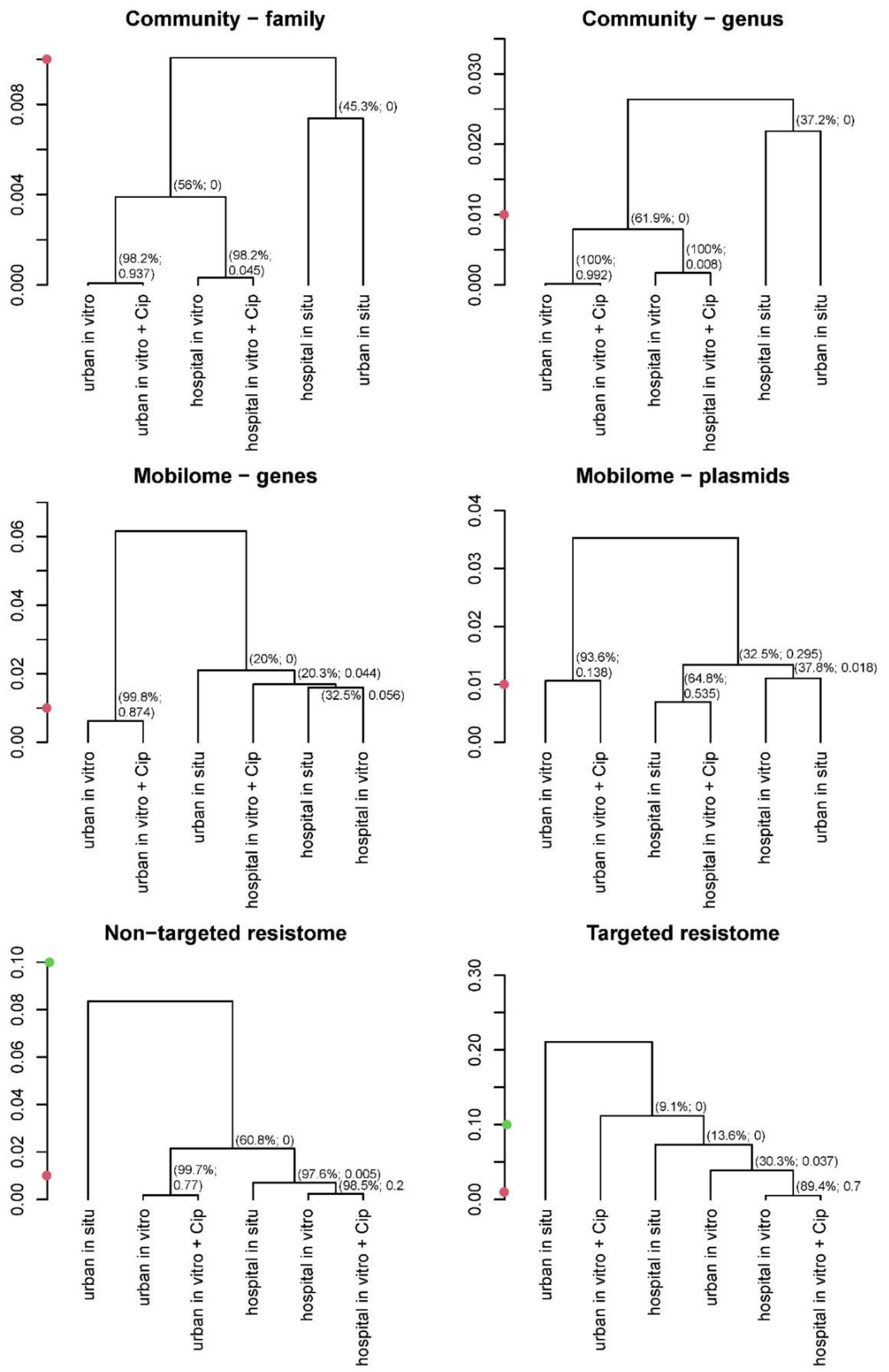


**Supplementary Figure 5:** UPGMA trees based on the proportion of Simpson diversity between samples. For each of the six multivariate features describing the biofilms, divergence between each pair of samples was assessed by calculating the proportion of the total Simpson diversity between samples. % Indicates the boot-strap robustness and the numeric values on the right correspond to conservative *P*- values, as described in methods. Red and green dots indicate the values 0.01 and 0.1, and they are devised to help the comparison between trees.

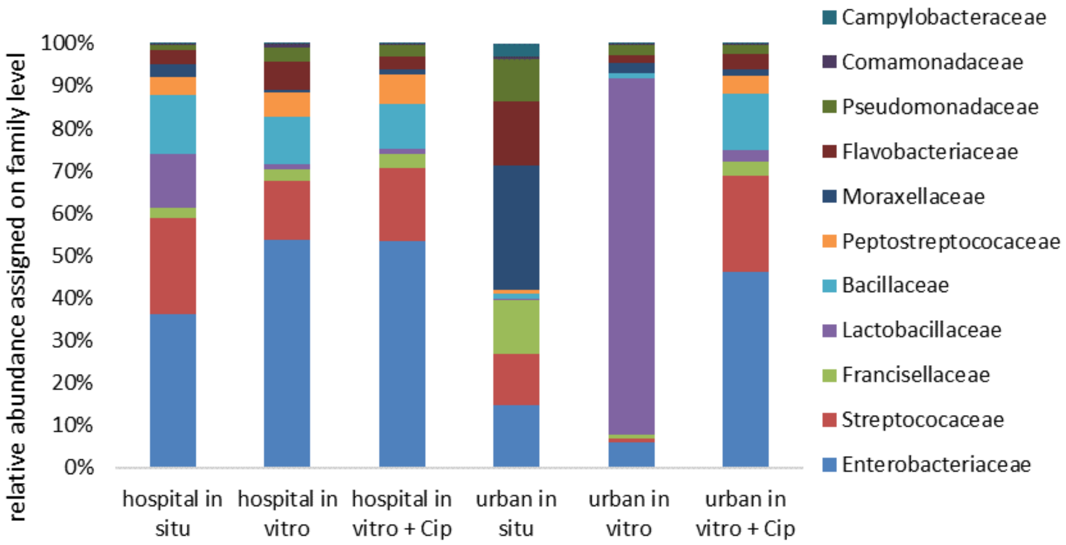


**Supplementary Figure 6:** Bacterial families identified by analysis of 16S rRNA reads after rRNA depletion *in vitro,* by kraken on the Silva database.

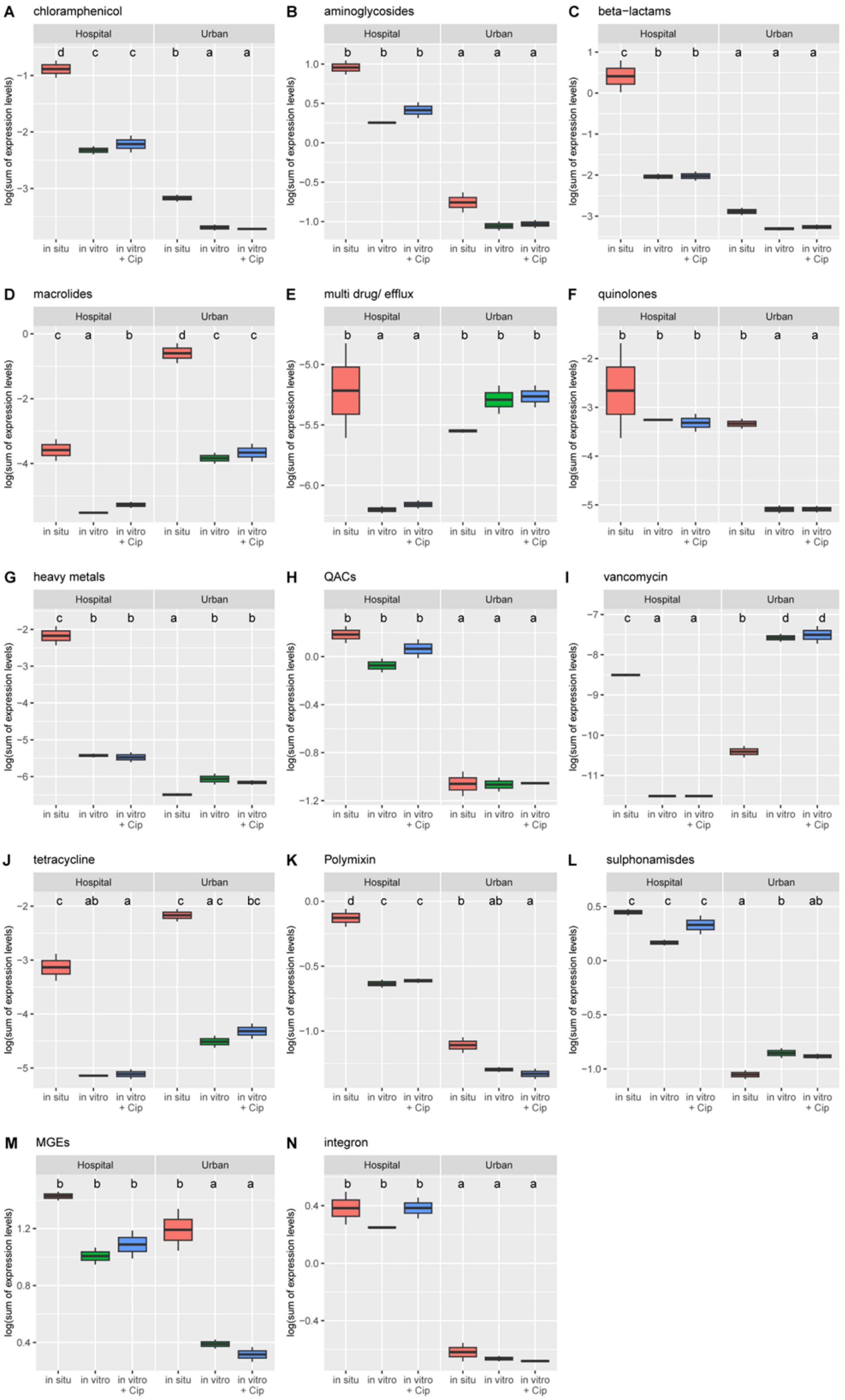


**Supplementary Figure 7: Comparisons of normalized abundances of the ARGs grouped into gene classes.** For each class of resistance genes, the boxplot shows the total normalized abundance per gene class and sample. The significance, (adjusted for multiple testing with the single step method) of the pairwise comparison is summarized by letters (compact letter display): samples that do not share a letter in common are significantly different of each other.

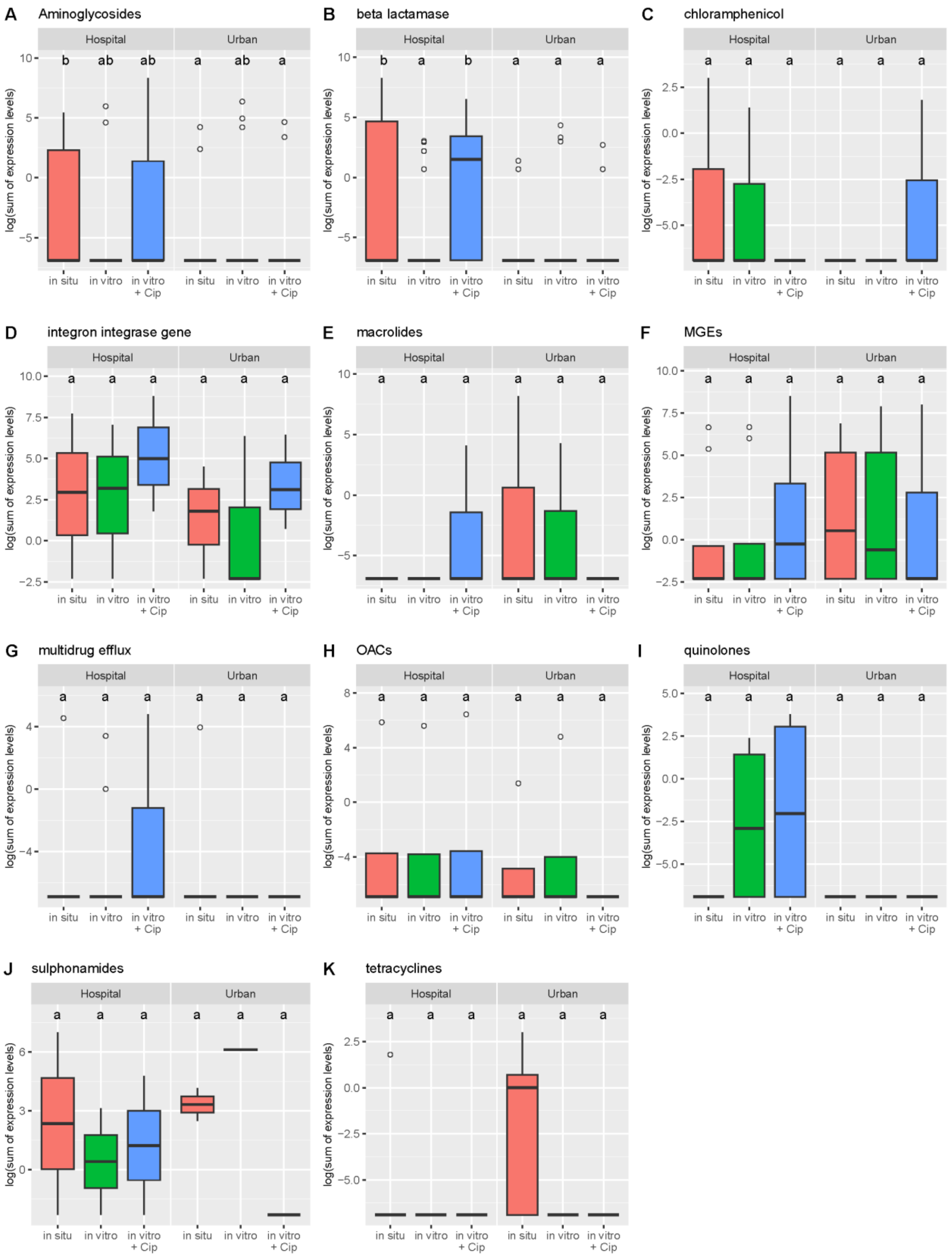


**Supplementary Figure 8: Comparisons of mean mapped mRNA reads corresponding to ARGs for the targeted resistome grouped into gene classes.** For each class of resistance genes, the boxplot shows the summed mean mapped mRNA reads per gene class and sample. The significance, (adjusted for multiple testing with the single step method) of the pairwise comparison is summarized by letters (compact letter display): samples that do not share a letter in common are significantly different of each other.

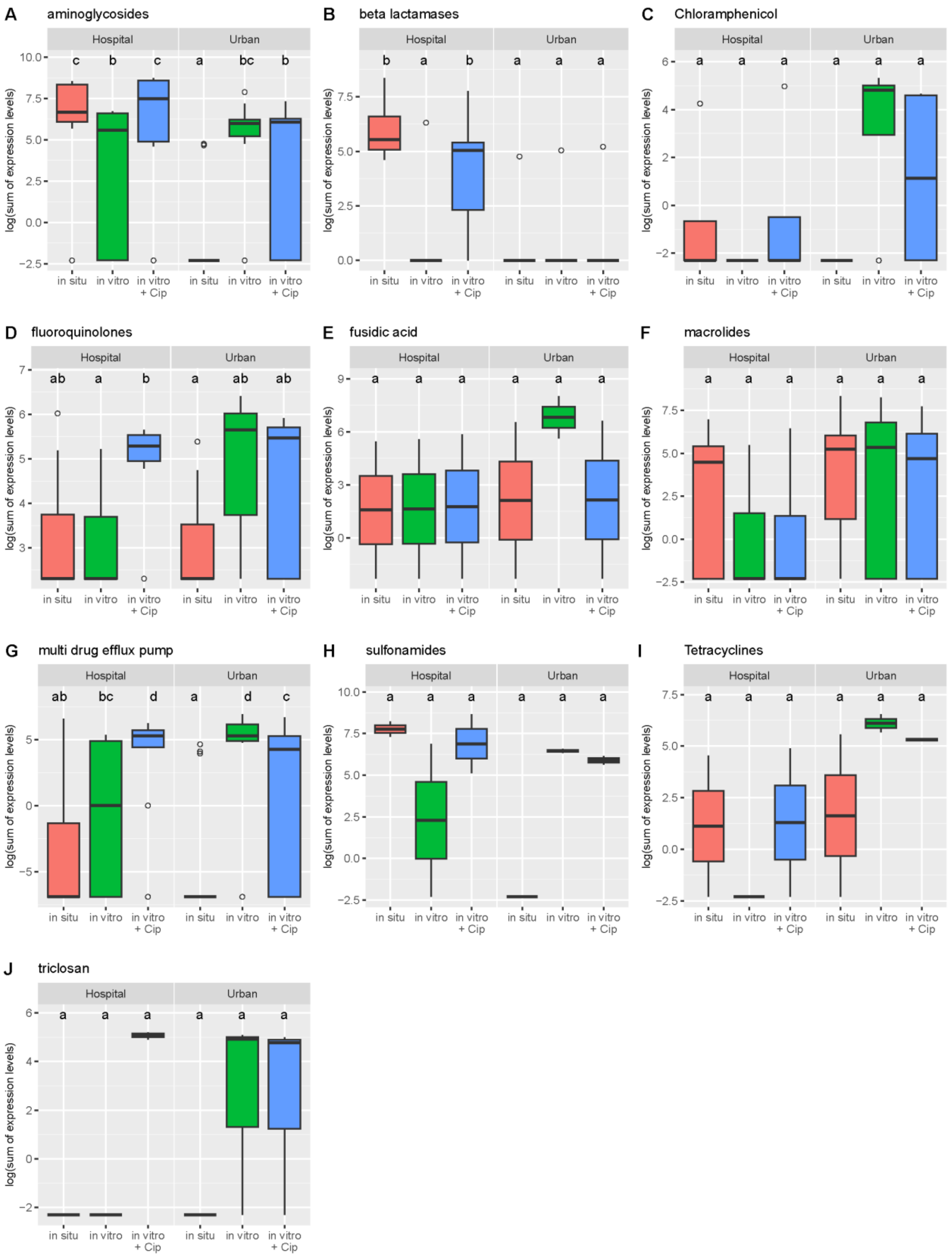


**Supplementary Figure 9: Comparisons of mean mapped mRNA reads corresponding to ARGs for the CARD resistome grouped into gene classes.** For each class of resistance genes, the boxplot shows the summed mean mapped mRNA reads per gene class and sample. The significance, (adjusted for multiple testing with the single step method) of the pairwise comparison is summarized by letters (compact letter display): samples that do not share a letter in common are significantly different of each other.

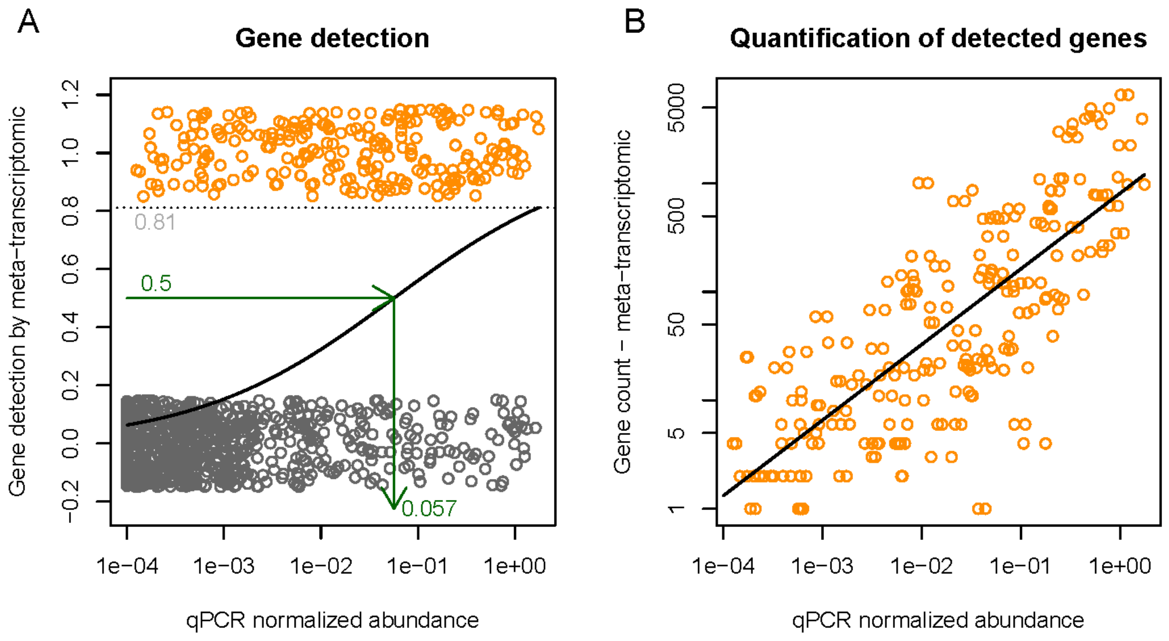


**Supplementary Figure 10: Correlation of the normalized abundance for ARGs detected by qPCR and their likelihood to be detected by our meta-transcriptomic pipeline.** Panel **A** analyzes the gene detection power of the meta-transcriptomic approach as a function of the genes normalized abundance obtained by qPCR. Noise has been added to the Y axis to make dots distinguishable. Black line indicates the binomial model prediction. Green arrows indicate the calculation of the detection threshold, and the dotted grey line give the maximal detection power we get for genes with the highest normalized abundance obtained by qPCR. Panel **B** shows the linear relation between the quantification by qPCR and meta-transcriptomic. Black line indicates the Poisson model prediction.

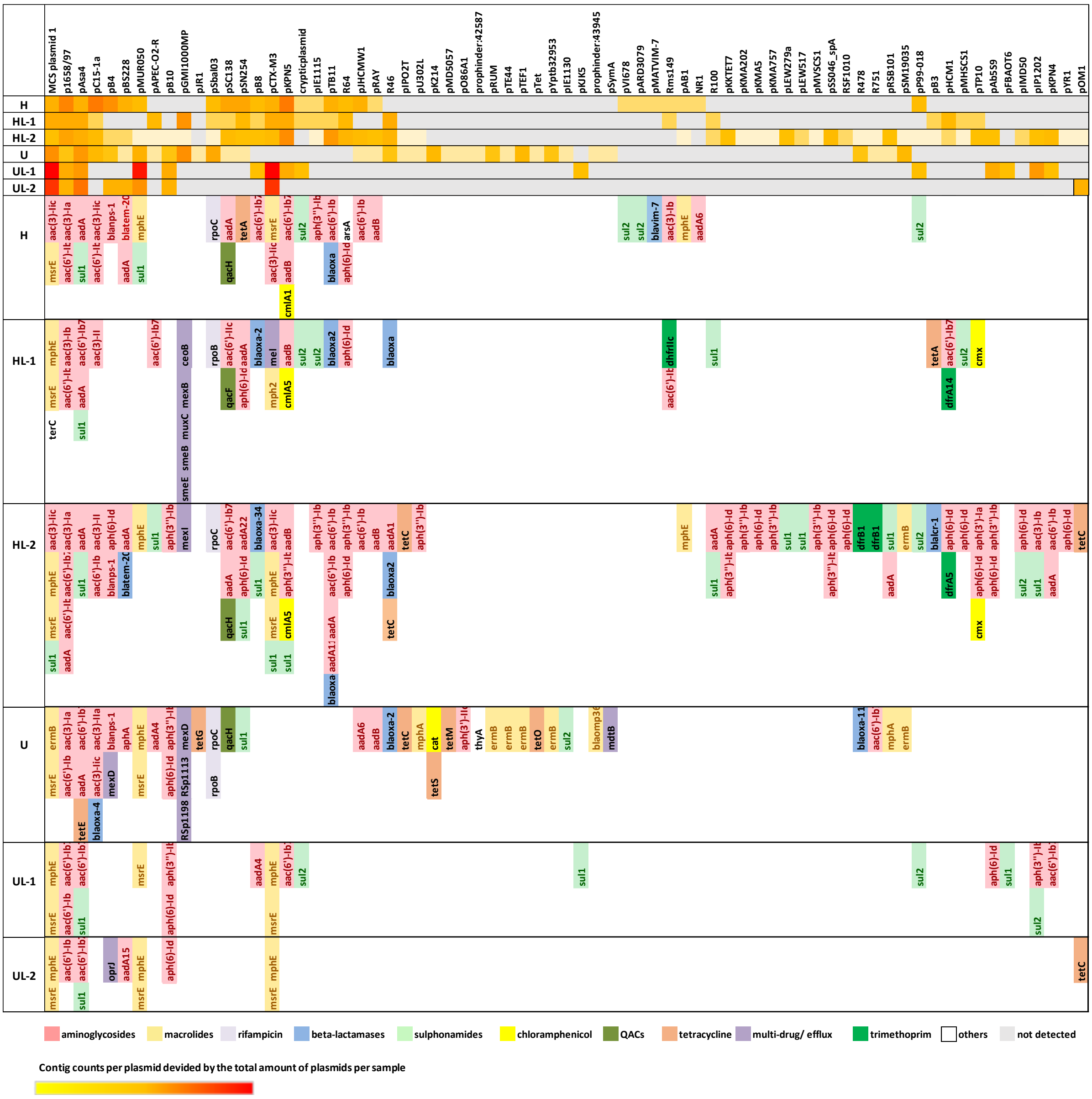


**Supplementary Figure 11: Active resistance mobilome:** Quantitative (heatmap) and qualitative (color coded table) representation of the active resistance mobilome in all biofilm samples. Counted and assigned MGEs (plasmids) are indicated in the heatmap for the respective samples. The results of the MGEs contig mapping on the CARD database, which revealed the actively expressed ARGs present on those MGEs is enlisted below, genes are color g=coded according to their resistance gene class. Samples: H = in situ hospital wastewater biofilm; HL-1 = in vitro hospital Wastewater biofilm; HL-2 in vitro hospital wastewater biofilm exposed to ciprofloxacin for 4 days. U = in situ urban wastewater biofilm; UL-1 = in vitro urban wastewater biofilm; UL-2 in vitro urban wastewater biofilm exposed to ciprofloxacin for 4 days.

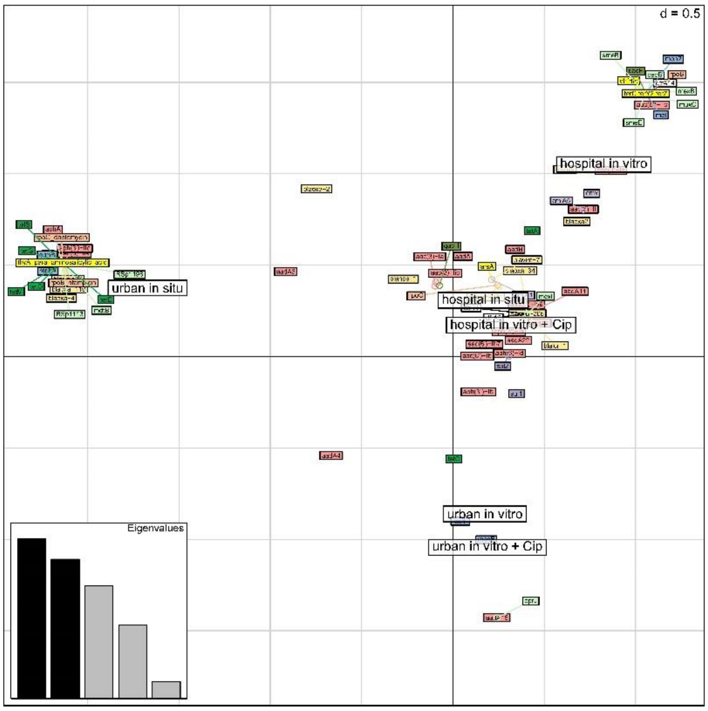


**Supplementary Figure 12:** Correspondence analysis (CA) to analyze the association between the class of the ARGs of the mobilome and the WW origin of the biofilm and their treatment (*in situ*, *in vitro*, with or without ciprofloxacin, CA; function ‘dudi.coa’ of the ade4 R package).
